# CITED2 is a Conserved Regulator of the Uterine-Placental Interface

**DOI:** 10.1101/2022.06.15.496287

**Authors:** Marija Kuna, Pramod Dhakal, Khursheed Iqbal, Esteban M. Dominguez, Lindsey N. Kent, Masanaga Muto, Ayelen Moreno-Irusta, Keisuke Kozai, Kaela M. Varberg, Hiroaki Okae, Takahiro Arima, Henry M. Sucov, Michael J. Soares

## Abstract

Establishment of the hemochorial uterine-placental interface requires exodus of trophoblast cells from the placenta and their transformative actions on the uterus, which represent processes critical for a successful pregnancy, but are poorly understood. We examined the involvement of CBP/p300-interacting transactivator with glutamic acid/aspartic acid-rich carboxyl terminal domain 2 (**CITED2**) in rat and human trophoblast cell development. The rat and human exhibit deep hemochorial placentation. CITED2 was distinctively expressed in the junctional zone and invasive trophoblast cells of the rat. Homozygous *Cited2* gene deletion resulted in placental and fetal growth restriction. Small *Cited2* null placentas were characterized by disruptions in the junctional zone, delays in intrauterine trophoblast cell invasion, and compromised plasticity. In the human placentation site, CITED2 was uniquely expressed in the extravillous trophoblast (**EVT**) cell column and importantly contributed to development of the EVT cell lineage. We conclude that CITED2 is a conserved regulator of deep hemochorial placentation.

**Significance Statement:** The process of establishing the uterine-placental interface is a poorly understood tissue re-engineering event that involves genetically foreign trophoblast cells breaching the immunologically secure uterus. When optimal, mother and fetus thrive, whereas failures represent the root cause of life-threatening diseases of pregnancy. CBP/p300-interacting transactivator with glutamic acid/aspartic acid-rich carboxyl terminal domain 2 (**CITED2**) is a transcriptional co-regulator with a conspicuous presence in trophoblast cell lineages infiltrating the uterine parenchyma. CITED2 helps coordinate the differentiation of rat and human trophoblast cells into invasive/extravillous trophoblast cells capable of transforming the uterus. These actions ensure requisite placental development and adaptations to physiological stressors. CITED2 exemplifies a conserved regulator of transcriptional events essential for establishing the uterine-placental interface.

## Introduction

The hemochorial placenta creates an environment essential for survival and development of the fetus (1–3). Several essential tasks are accomplished by the placenta. Trophoblast, the parenchymal cell lineage of the placenta, specializes into cell types facilitating the flow of nutrients into the placenta and their transfer to the fetus (3, 4). Fundamental to this process is the differentiation of trophoblast cells with the capacity to enter and transform uterine tissue proximal to the developing placenta, and restructure uterine vasculature (3, 4). Intrauterine trophoblast cell invasion and trophoblast cell-guided uterine transformation are highly developed in the human and the rat, unlike the mouse (5, 6). In the human, these cells are referred to as extravillous trophoblast (**EVT**) cells and the generic term, invasive trophoblast cells, is used to identify these cells in the rat. Failure of trophoblast cell-directed uterine transformation has negative consequences for mother and fetus (4). Invasive trophoblast cell progenitors arise from structures designated as the EVT cell column and the junctional zone in the human and rat, respectively (7). The EVT cell column is a well-defined structure containing a stem/proliferative population of trophoblast cells (cytotrophoblast) situated at the base of the column with a linear progression of EVT progenitor cells located within the core of the column proceeding to various stages of maturing EVT cells positioned at the distal region of the column (4). In contrast, the junctional zone is more complex, giving rise to endocrine cells (trophoblast giant cells and spongiotrophoblast cells), energy reservoirs (glycogen trophoblast cells), and invasive trophoblast progenitor cells (6, 8). Understanding cellular decision-making within the EVT cell column and junctional zone provides insights into the development of the invasive trophoblast cell lineage.

CBP/p300 interacting transactivator, with Glu/Asp-rich carboxy terminal domain, 2 (**CITED2**) is a transcriptional co-regulator possessing the capacity to modulate interactions between DNA binding proteins and histone modifying enzymes, specifically transcription factor-CREB binding protein (**CREBBP or CBP**)/EIA binding protein p300 (**EP300**) interactions (9, 10). CBP and EP300 possess histone 3 lysine 27 (**H3K27**) acetyl transferase activity (11, 12) and have been implicated in trophoblast cell differentiation and their dysregulation linked to diseases of the placenta, including preeclampsia and intrauterine growth restriction (13, 14). The outcome of CITED2 actions is transcription factor specific. In some cases, CITED2 interferes with transcription factor-CBP/EP300 interactions and inhibits gene expression (e.g. hypoxia inducible factor, **HIF**) (15), while in other cases, CITED2 facilitates transcription factor-CBP/EP300 recruitment and activates gene expression (e.g., Activator protein 2 family, **TFAP2**) (16). Both HIF and TFAP2C (also called **AP-2**γ) have essential roles in placentation (17–21). The connections between CITED2 and these molecular targets place CITED2 at key positions in the regulatory network controlling trophoblast cell development. In fact, mutagenesis of the mouse *Cited2* locus results in placental malformation (22, 23), along with a range of other embryonic defects, including prenatal lethality (15, 16, 24). Furthermore, CITED2 is prominently upregulated during rat trophoblast cell differentiation (25, 26), suggesting it may directly facilitate placental development.

In this report, we explore the involvement of CITED2 in the regulation of trophoblast cell development and deep placentation using genetically manipulated rat models and trophoblast stem (**TS**) cells. *CITED2* is expressed in the junctional zone and invasive trophoblast cells of the rat placentation site. Disruption of *Cited2* results in compromised growth of the junctional zone, abnormalities in the invasive trophoblast cell lineage, dysregulation of TS cell differentiation, and abnormalities in adaptive responses to hypoxia and immune challenges. We also describe prominent phenotypic differences between mice and rats possessing *Cited2* null mutations. Importantly, we show that CITED2 is a conserved regulator of invasive trophoblast/EVT cell lineage decisions.

## Results

### Cited2 expression within the rat placentation site

We started our investigation of the involvement of CITED2 in deep placentation by examining *Cited2* expression in the rat. The rat placentation site is organized into three well-defined compartments (labyrinth zone, junctional zone, and uterine-placental interface) that can be enriched by dissection (**Fig. 1A**). The labyrinth zone is situated at the placental-fetal interface adjacent to the junctional zone, which borders the uterine parenchyma. As gestation progresses, invasive trophoblast cells detach from the junctional zone and infiltrate the uterine parenchyma, establishing a structure we define as the uterine-placental interface, which has also been called the metrial gland (6, 7). Reverse transcription-quantitative polymerase chain reaction (**RT-qPCR**) measurements demonstrated abundant expression of *Cited2* transcripts in the junctional zone, which was far greater than any other tissue analyzed (**Fig. 1B**). Expression of *Cited2* increased in the junctional zone and uterine-placental interface as gestation progressed (**Fig. 1C and D**). Localization of *Cited2* transcripts confirmed their presence in the junctional zone and within invasive trophoblast cells of the uterine-placental interface (**Fig. 1E**). The latter was demonstrated by co-localization of *Cited2* and *Prl7b1* transcripts. *Prl7b1* is an established marker of the invasive trophoblast cell lineage of the rat (27). Thus, *Cited2* is present in compartments of the placentation site critical to the derivation (junctional zone) and functioning of invasive trophoblast cells (uterine-placental interface).

**Figure 1.**
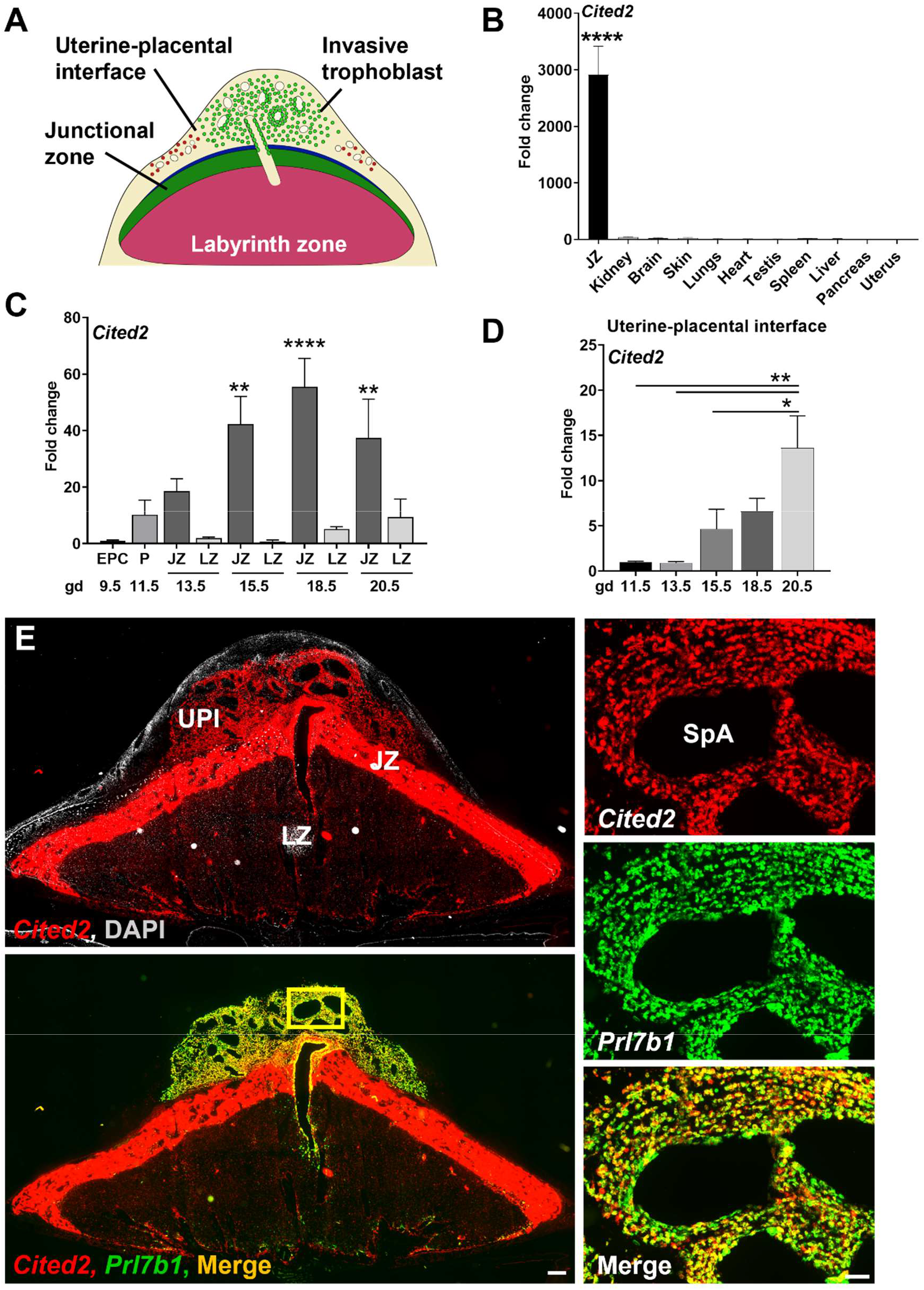
*Cited2* expression in the placenta and uterine-placental interface during gestation in the rat. **A**. Schematic showing the late gestation rat placentation site. Invaded trophoblast cells are depicted in green. **B**. Relative expression of *Cited2* transcript in postnatal day 1 (**PND1**) rat neonatal tissues and gestation day (**gd**) 14.5 junctional zone (**JZ**) tissue. **C**. Relative expression of *Cited2* transcripts in the ectoplacental cone (**EPC**), whole placenta (**P**), JZ, and labyrinth zone (**LZ**) of the rat placenta during gestation. Values depicted were normalized to gd 9.5 EPC samples. **D**. Relative expression of *Cited2* transcripts within the uterine-placental interface during gestation. **E**. *In situ* hybridization showing *Cited2* transcript distribution (top left) and *Cited2* and *Prl7b1* (invasive trophoblast marker) transcript co-localization in rat gd 18.5 placentation site (bottom left). Higher magnification images of the area outlined by a yellow rectangle (bottom left) are shown to the right. Scale bar=500 μm (left panels), scale bar=100 μm (right panels). Uterine-placental interface (**UPI**), spiral artery (**SpA**). The histograms presented in panels **B, C**, and **D** represent means ± SEM, n=5-10, 3-6 pregnancies. One-way ANOVA, Tukey”s post hoc test, * p < 0.05, ** p < 0.01, **** p<0.0001.

### In vivo analysis following Cited2 disruption

A global *Cited2* mutant rat model was generated using *CRISPR*/Cas9 mediated genome-editing. Two guide RNAs were used to generate a 1477 bp nucleotide deletion, which removed the entire coding region of *Cited2* (***SI Appendix* Fig. S1A and B**). The *Cited2* mutant allele was successfully transferred through the germline. Both male and female rats heterozygous for the *Cited2* mutation were viable and fertile. The absence of CITED2 protein in the placenta of homozygous mutants confirmed the gene disruption (***SI Appendix* Fig. S1C**). Since *Cited2* null pups were not observed at weaning from heterozygous x heterozygous breeding, we hypothesized that *Cited2* null rats died in utero, as observed for *Cited2* mutant mice (15, 16, 24) or soon after birth. In contrast to the mouse, *Cited2* null mutant rats survived prenatal development and instead, died postnatally, within a few h of extrauterine life (***SI Appendix* Fig. S1D**). This fundamental difference prompted a more detailed comparison of the effects of *Cited2* disruption in the rat versus the mouse. We obtained a well-characterized *Cited2* mutant mouse model, which also possessed a deletion of the entire coding sequence (***SI Appendix* Fig. S1E**) (15). The *Cited2* mutation was transferred to an outbred CD1 mouse genetic background following backcrossing for >10 generations. It has been reported that disruption of the *Cited2* gene in the mouse results in fetal growth restriction and a range of developmental anomalies, including: i) cardiac abnormalities; ii) arrested lung development; iii) absence of adrenal glands, and iv) neural tube defects resulting in exencephaly (15, 16, 24, 28, 29). Fetal rats possessing homozygous *Cited2* mutations exhibited heart and lung abnormalities (***SI Appendix* Fig. S2A and B**) as previously reported for the mouse (15, 16, 28, 29). At embryonic day (**E**) 15.5, all *Cited2* null rat hearts examined possessed ventral septal defects and double outlet right ventricle and half showed a retroesophageal right subclavian artery (***SI Appendix* Fig. S2A**) (30). Connections between abnormal placentation and fetal heart defects have been previously described (31–33). Postnatal day 1 lung development in *Cited2* homozygous mutant rats failed to progress and was arrested at the canicular stage (***SI Appendix* Fig. S2B**) (34). Failures in heart and lung development are probable causes of death of *Cited2* nulls on the first day of extrauterine life. In contrast to the mouse, disruption of the *Cited2* gene in the rat showed no detectable adverse effects on adrenal gland or neural tube development (***SI Appendix* Fig. S2C-G**). Thus, similarities and prominent differences exist in the phenotypes of rats versus mice with *Cited2* null mutations.

### CITED2 deficiency leads to placental growth restriction

In the rat, global *Cited2* deficiency resulted in placental and fetal growth restriction starting at gestation day (**gd**) 14.5 and persisted through the end of gestation (**Fig. 2A**). Similar placental and fetal growth deficits were observed for the *Cited2* null mouse (***SI Appendix* Fig. S3**), as previously reported (22). Both wild type and *Cited2* null rat placentas were organized into well-defined labyrinth and junctional zone compartments; however, each compartment was significantly smaller in *Cited2* null placentation sites (**Fig. 2B and C**). Genotype-dependent differences were not noted in the expression of proliferation or apoptotic markers in the gd 13.5 and 14.5 rat junctional zone (***SI Appendix* Fig. S4**).

**Figure 2.**
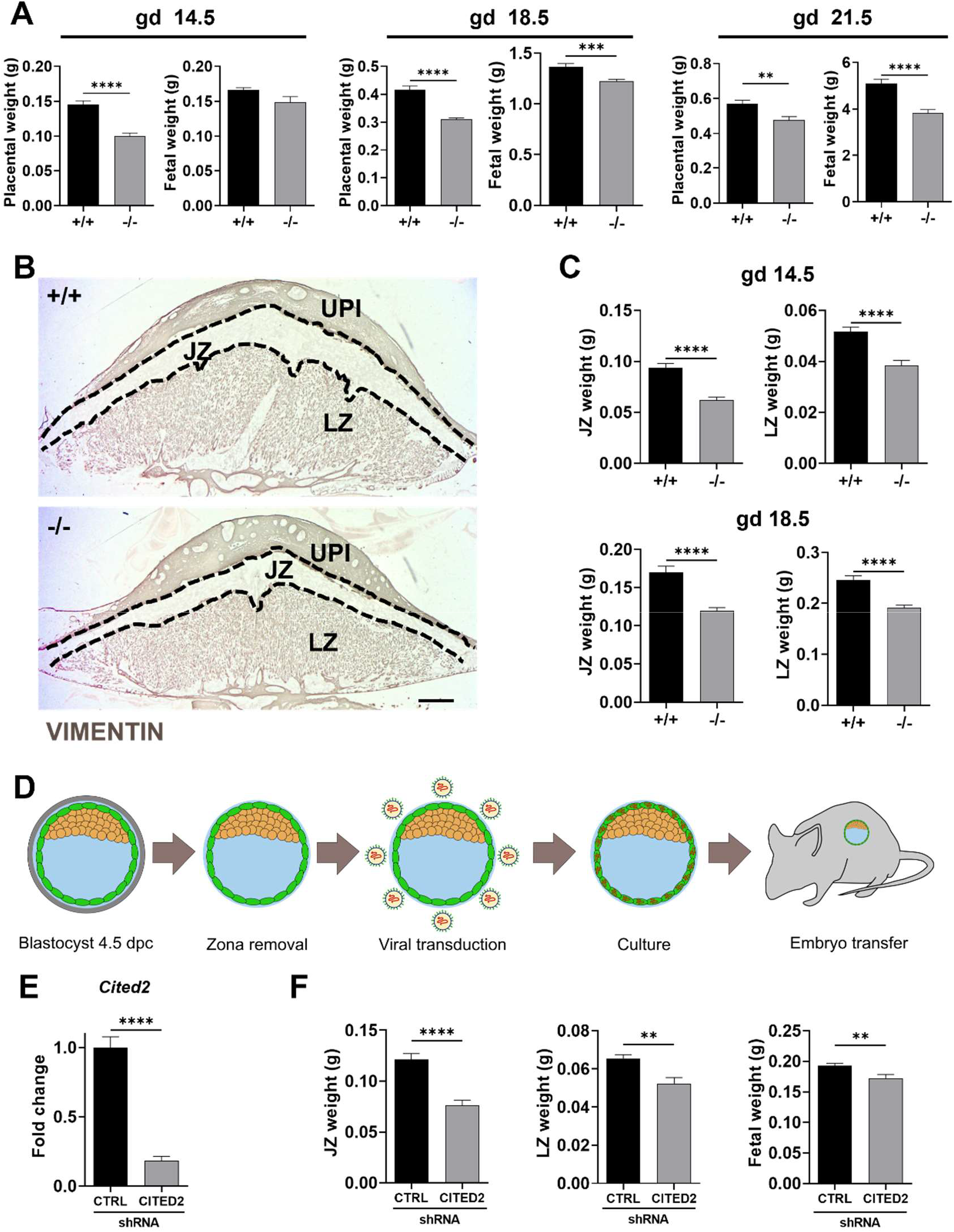
The CITED2 deficient rat placenta is growth restricted. **A**. Placental and fetal weights from *Cited2*+/-x *Cited2*+/-breeding for the rat. **B**. Immunohistological analysis of vimentin in gestation day (**gd**) 18.5 wild type (+/+) and null (-/-) placentas from *Cited2+/-* x *Cited2+/-* breeding. Scale bar=1000 μm. **UPI**, uterine-placental interface, **JZ** junctional zone, **LZ** labyrinth zone. **C**. JZ and LZ weights from *Cited2+/-* x *Cited2+/-* breeding on gd 14.5 and gd 18.5. Values represent mean ± SEM, n=12-35, unpaired t-test, **p<0.01, ***p<0.001, **** p<0.0001. **D**. Simplified schematic depicting the strategy for achieving trophoblast specific CITED2 knockdown in vivo. **E**. Relative expression of *Cited2* transcripts in control (**CTRL**) and CITED2 shRNA-exposed gd 14.5 JZ tissue. **F**. JZ, LZ, and fetal weights from control and gd 14.5 trophoblast specific *Cited2* knockdown. Control (**CTRL**) and *Cited2* shRNA mediated knockdown. Shown are mean values ± SEM, n=12-20, unpaired t-test, **p<0.01, **** p<000.1.

Using trophoblast-specific lentiviral delivery (35) of *Cited2*-specific short hairpin RNAs (**shRNA**) (26), we determined that the effects of CITED2 on placental size and function were trophoblast-specific (**Fig. 2D-F**). *Cited2* shRNAs were transduced into trophectoderm of blastocysts. Transduced blastocysts were transferred into pseudopregnant female rats, and placental and fetal size evaluated at gd 14.5. Junctional zone, labyrinth zone, and fetal weights were significantly smaller in *Cited2* shRNA transduced trophoblast versus control shRNA transduced trophoblast (**Fig. 2F**).

The effects of CITED2 deficiency on the junctional zone could be viewed as a cell autonomous action, whereas growth defects in the labyrinth zone as a non-cell-autonomous action, potentially arising from deficits in the junctional zone or its derivatives, including invasive trophoblast cells and their actions in transforming the uterine parenchyma.

### CITED2 deficiency affects gene regulatory networks in the junctional zone

We next examined the potential cell-autonomous actions of CITED2 on junctional zone development. CITED2 is a known transcriptional co-regulator (9, 10), which prompted an examination of CITED2 deficiencies on the junctional zone gene regulatory network. RNA sequencing (**RNA-seq**) was performed on gd 14.5 wild type and *Cited2* null junctional zone tissues. A total of 203 differentially regulated transcripts were identified in the RNA-seq analysis, which included the downregulation of 160 transcripts and upregulation of 43 transcripts in CITED2 deficient junctional zone tissue (***SI Appendix* Fig. S5A-C, Dataset S1**). Among the downregulated transcripts were transcripts known to be prominently expressed in the junctional zone (e.g., *Mmp9, Igf2, Cyp11a1, Prl3d4, Prl8a9*). Surprisingly, an assortment of known interferon-responsive transcripts was upregulated in CITED2 deficient junctional zone tissues (e.g., *Ifi27l2b, Isg15, Oas1f, Ifitm3*; ***SI Appendix* Fig. S5C**). Pathway analysis supported roles for CITED2 in the regulation of cell migration and immune effector processes (***SI Appendix* Fig. S5D**). Rat TS cells represent an excellent model for junctional zone development (36). *Cited2* transcript levels were dramatically upregulated following rat TS cell differentiation (***SI Appendix* Fig. S6A**). Consequently, we derived rat TS cells from CITED2 deficient blastocysts and compared their behavior to wild type rat TS cells (***SI Appendix* Fig. S6B-E**). Morphologies of wild type and *Cited2* null rat TS cells were similar in the stem state and following differentiation (***SI Appendix* Fig. S6C**). CITED2 deficient TS cells grew slower than wild type rat TS cells (***SI Appendix* Fig. S6D**) and exhibited dysregulated gene expression as determined by RT-qPCR (***SI Appendix* Fig. S6E**). Interestingly, two transcripts known to regulate placental development, including the invasive trophoblast cell lineage, *Mmp9* and *Igf2* (37, 38), were similarly downregulated in CITED2 deficient rat TS cells (***SI Appendix* Fig. S6E**).

### CITED2 deficiency and invasive trophoblast cell development

Trophoblast cells invade deep into the rat uterine parenchyma (5, 39). These intrauterine invasive trophoblast cells express *Cited2* (**Fig. 1E**), implicating CITED2 as a potential regulator of the development and/or function of the invasive trophoblast cell lineage. Early endovascular trophoblast cell invasion into the decidua at gd 13.5 was limited in *Cited2* nulls in comparison to wild type placentation sites; however, as gestation progressed differences in the extent of intrauterine trophoblast invasion was not evident between *Cited2* nulls and wild type placentation sites (**Fig. 3A-D**). The invasive trophoblast cell developmental delay characteristic of *Cited2* null placentation sites was evident at gd 15.5, as visualized by in situ hybridization for *Ceacam9* and *Prl7b1* (**Fig. 3C**). Prominent phenotypic differences in wild type and CITED2 deficient invasive trophoblast cells emerged from single cell RNA-seq (**scRNA-seq**) of the gd 18.5 uterine-placental interface (**Fig. 4, *SI Appendix* Fig. S7A-E, Table S1, Datasets S2-S5**). The uterine-placental interface from gd 18.5 was selected to obtain sufficient number of invasive trophoblast cells for analysis. Uniform manifold approximation and projection (**UMAP**) profiles of the uterine-placental interface were similar to previously published UMAP profiles for the rat uterine-placental interface (40). *Cited2* expression was enriched in the invasive trophoblast cell cluster (**Fig. 4B**). Disruption of CITED2 did not prevent the development of invasive trophoblast cells but altered their phenotype and their numbers (**Fig. 4C-E**). Among the differentially expressed invasive trophoblast cell transcript signatures sensitive to CITED2 was signaling by Rho GTPases (**Fig. 4F**), which is fundamental to the regulation of cell migration and invasion (41). Transcripts encoding cell adhesion molecules (CEACAM9, NCAM1), proteins promoting cell migration (CCDC88A), ligands targeting the vasculature (VEGFA, NPPB), and interferon-responsive proteins (IFITM3, IFI27L2B, IFI27) were differentially expressed. Interestingly, *Doxl1* was downregulated in the absence of CITED2. DOXL1 is a paralog of AOC1, which encodes a diamine oxidase responsible for the oxidation of polyamines. AOC1 is prominently expressed in EVT cells and is dysregulated in disorders such as preeclampsia (42). Collectively, the findings indicate that CITED2 regulates the invasive trophoblast cell lineage.

**Figure 3.**
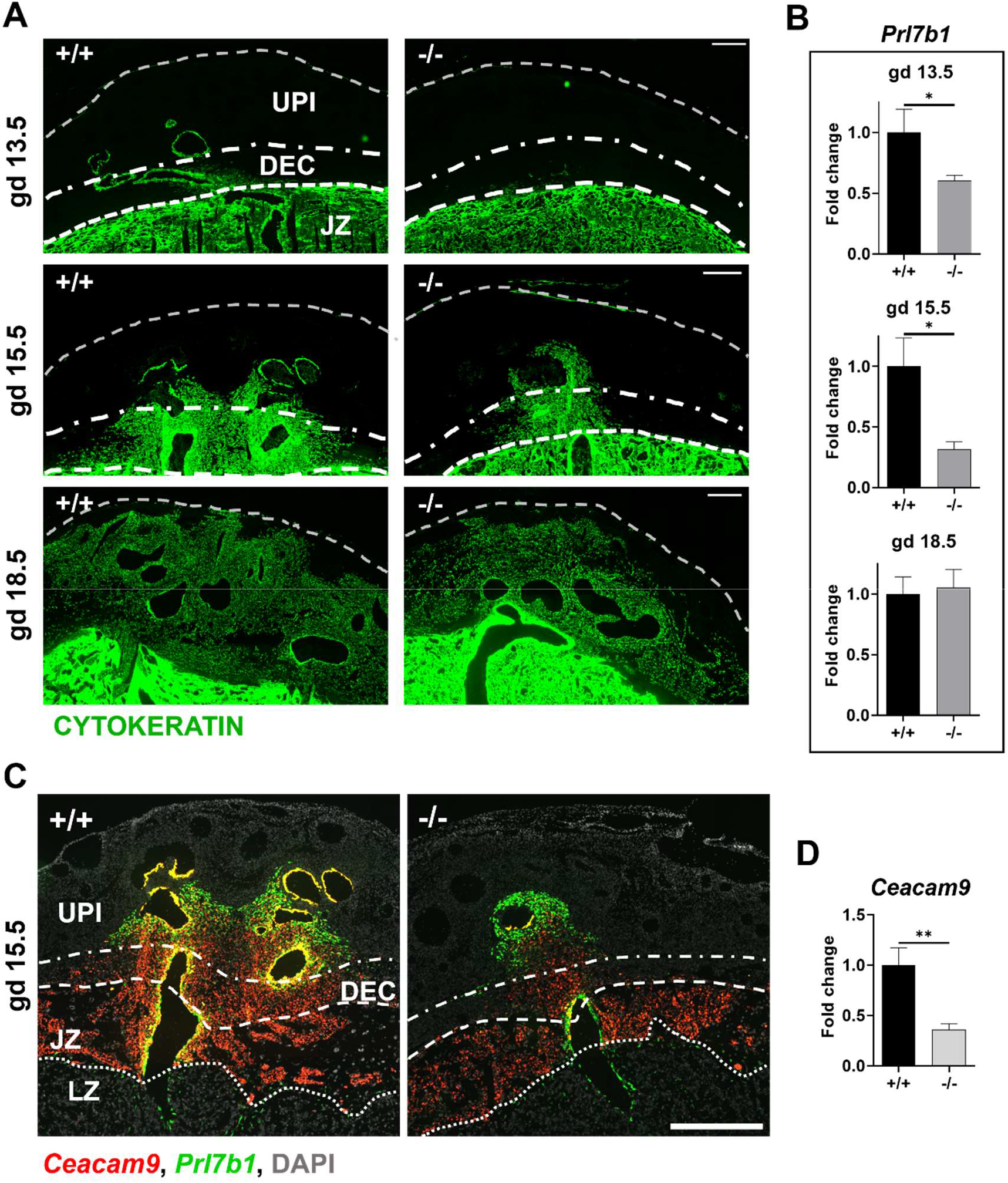
Intrauterine trophoblast cell invasion is delayed in *Cited2* null rat placentation sites. **A**. Representative images of wild type (+/+) and *Cited2* null (-/-) rat gestation day (**gd**) 13.5, 15.5 and 18.5 placentation sites immunostained for cytokeratin (green). The cytokeratin immunostain is specific to invasive trophoblast cells that have entered the uterine parenchyma. Scale bars=500 μm. **B**. Relative expression of *Prl7b1* transcripts (invasive trophoblast cell marker) from wild type (+/+) and *Cited2* null (-/-) gd 13.5 decidual tissue and uterine placental interface tissue at gd 15.5 and 18.5 measured by RT-qPCR. Graphs depict mean values ± SEM, n=11-24, unpaired t-test, *p<0.05. **C**. *In situ* hybridization showing *Ceacam9* (invasive trophoblast cell marker, red) and *Prl7b1* (invasive trophoblast marker, green) transcript localization in gd 15.5 wild type (+/+) and *Cited2* null (-/-) rat placentation sites, scale bar=1000 μm. **D**. Relative expression of *Ceacam9* transcripts (invasive trophoblast cell marker) in gd 15.5 uterine placental interface tissue. Shown are mean values ± SEM, n=12-14, unpaired t-test, **p<0.01. Uterine placental interface (**UPI**), decidua (**DEC**), junctional zone (**JZ**), labyrinth zone (**LZ**).

**Figure 4.**
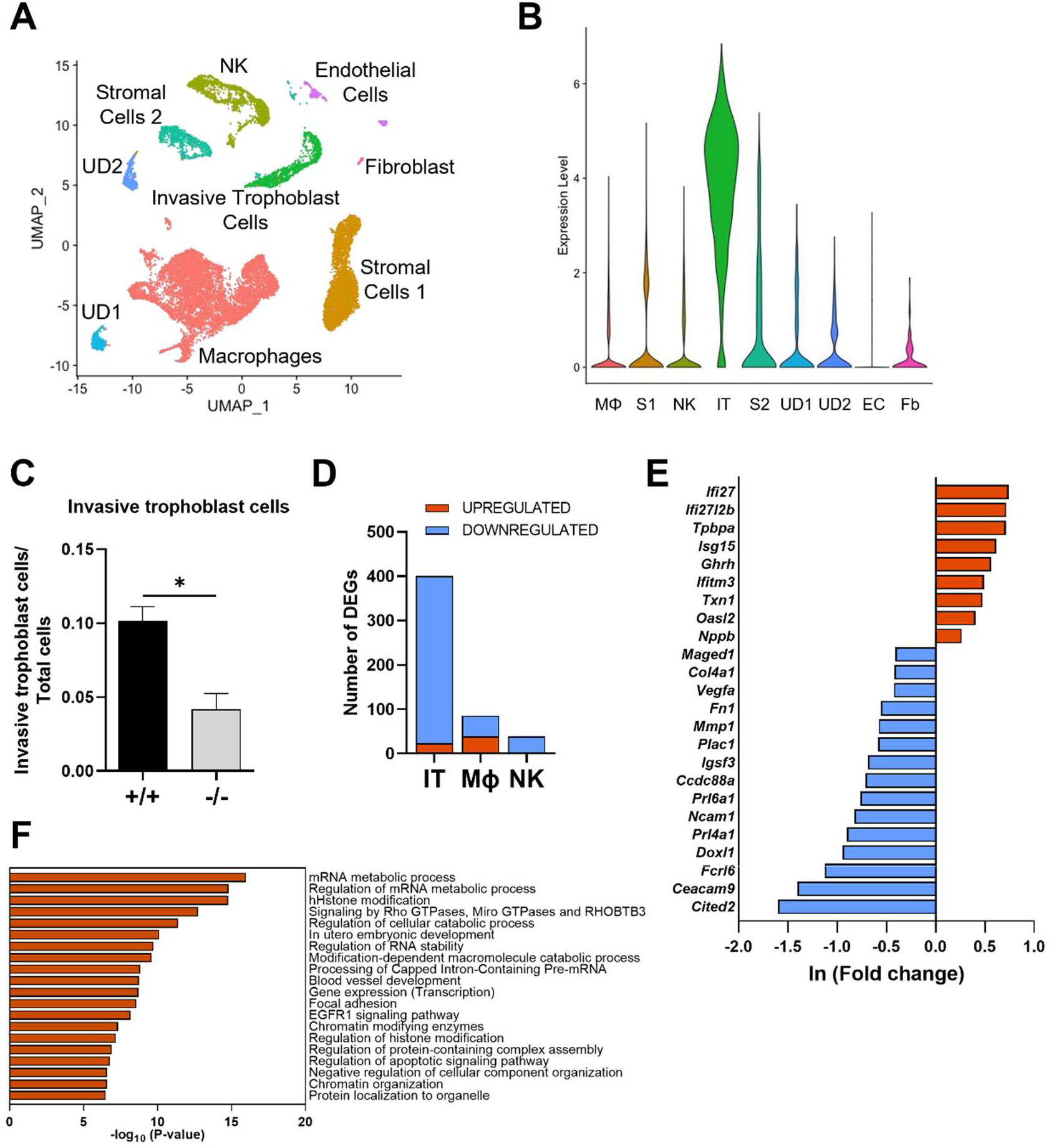
CITED2 deficiency affects the rat invasive trophoblast cell phenotype. Single cell-RNA sequencing was performed on wild type (+/+) and *Cited2* null (-/-) gestation day (**gd**) 18.5 uterine-placental interface tissue samples. **A**. UMAP plot showing cell clustering in wild type (+/+) and *Cited2* null (-/-) gd 18.5 uterine-placental interface tissue. **UD1**, undefined cell cluster 1; **UD2**, undefined cell cluster 2. **B**. Violin plot showing expression of *Cited2* in each cell cluster. Cell clusters: macrophages (**MΦ**) stromal 1 (**S1**), natural killer (**NK**) cells, invasive trophoblast (**IT**) cells, stromal 2 (**S2**) cells, UD1, UD2, endothelial cells (**EC**), and fibroblasts (**Fb**). **C**. Ratio of invasive trophoblast cells per total number of cells analyzed. Graph represents mean values ± SEM, n=3, unpaired t-test, *p<0.05. **D**. Number of differentially expressed genes (**DEGs**) for IT, **MΦ**, and **NK** cells from wild type (+/+) versus *Cited2* null (-/-) uterine-placental interface tissue. **E**. Bar plot showing select DEGs in the invasive trophoblast cell cluster (upregulated shown in red; downregulated shown in blue). **F**. Gene Ontology enriched terms for DEGs from the invasive trophoblast cell cluster.

### CITED2 and placentation site adaptations to physiological stressors

Placentation sites possess the capacity to adapt to exposure to physiological stressors (3, 7). Hypoxia and polyinosinic:polycytidylic acid (**polyI:C**), a viral mimic, can elicit placental adaptations (43, 44). Hypoxia exposure elicits a prominent increase in endovascular trophoblast cell invasion (43), whereas polyI:C can disrupt placental and fetal development (44). CITED2 is involved in the regulation of adaptations elicited by exposure to hypoxia and inflammation (15, 45, 46). Therefore, we investigated responses of wild type versus CITED2 deficient placentation sites to physiological stressors.

Maternal hypoxia exposure from gd 6.5 to 13.5 was sufficient to overcome the delay in endovascular trophoblast cell invasion observed in CITED2 deficient placentation sites (**Fig. 5A-C**). Additionally, when exposed to hypoxia CITED2 deficient conceptus sites showed an increased resorption rate compared to their wild type littermates, indicating that without CITED2 the adaptations to hypoxia are compromised (**Fig. 5D**).

**Figure 5.**
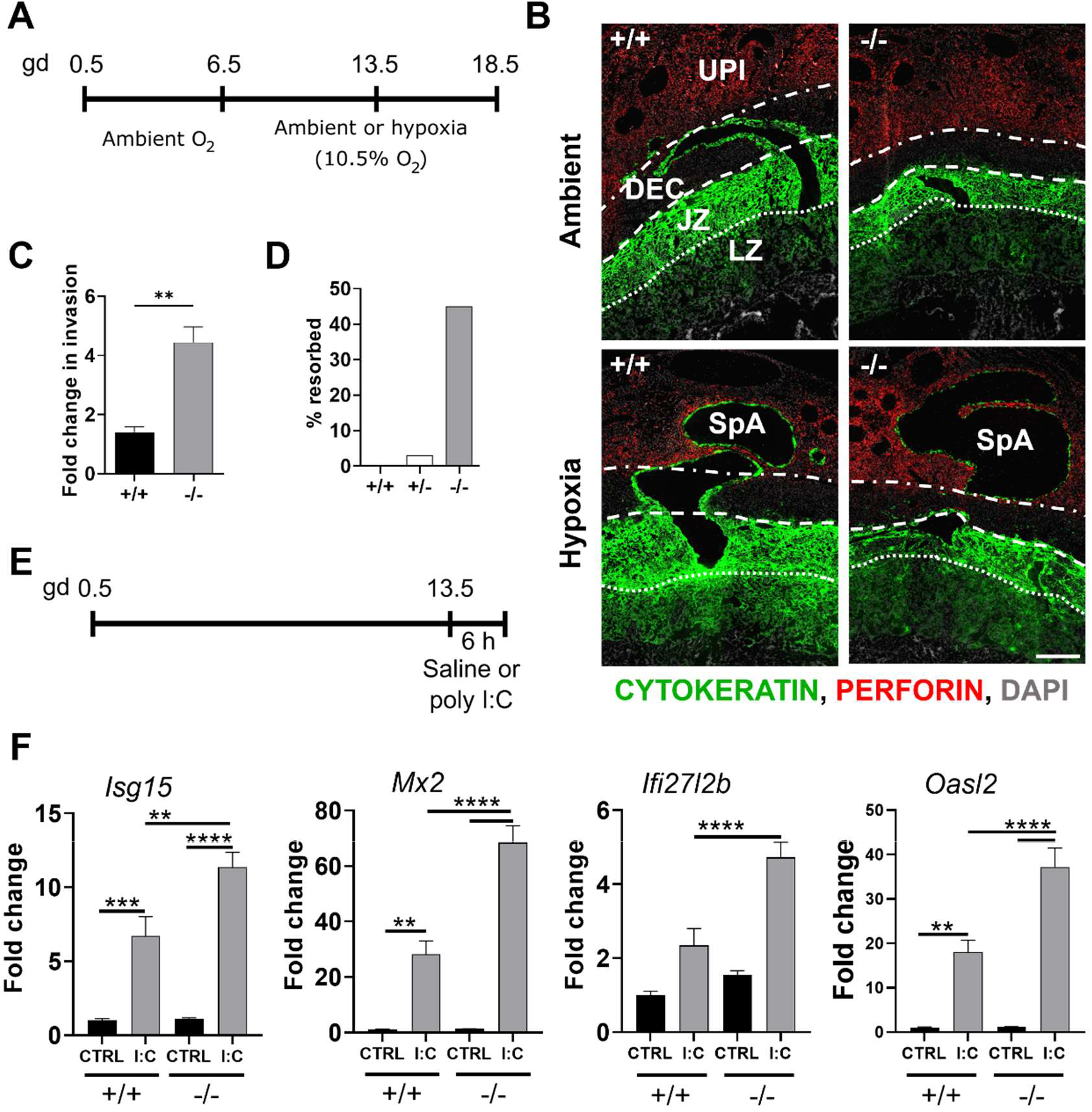
CITED2 modulates rat placental responses to physiological stressors. **A**. Schematic of the experimental timeline for hypoxia exposure. **B**. Representative images of wild type (+/+) and *Cited2* null (-/-) rat gestation day (**gd**) 13.5 placentation sites exposed to ambient or hypoxic (10.5% oxygen) conditions. Sections were immunostained for cytokeratin (green), perforin (red), and DAPI (blue). Scale bar=500 μm. Uterine placental interface (**UPI**), decidua (**DEC**), junctional zone (**JZ**), labyrinth zone (**LZ**), and spiral artery (**SpA**). **C**. Depth of gd 13.5 endovascular trophoblast invasion was quantified, and fold changes calculated for hypoxic relative to ambient conditions, n=5-9. **D**. Resorption rate assessed on gd 18.5 for individual genotypes from *Cited2*+/-females bred to *Cited2*+/-males and exposed to ambient of hypoxic conditions, n=20-33. **E**. Schematic of the experimental timeline for polyinosinic:polycytidylic acid (**polyI:C**) treatment, **F**. Relative expression of *Isg15, Mx2, Ifi27I2b*, and *Oasl2* in junctional zone tissue from control (saline treated; **CTRL**) and polyI:C exposed (**I:C**) wild type (+/+) and *Cited2* null (-/-) placentas; n=8-17. Shown are mean values ± SEM, one-way analysis of variance, Tukey”s post-hoc test. *p<0.05, **p <0.01, ***p <0.001, ****p <0.0001.

CITED2 deficiency was associated with upregulated expression of interferon-responsive transcripts in the junctional zone and invasive trophoblast cells (**Fig. 4E, *SI Appendix* Fig. S5, Dataset S1**), which implied that CITED2 could be involved in regulating responses to a viral challenge. We first determined the efficacy of a polyI:C challenge. PolyI:C treatment of pregnant rats resulted in significant increases in inflammatory transcript expression in the spleen and uterine-placental interface (***SI Appendix* Fig. S8**). We then measured interferon-responsive transcripts in wild type and *Cited2* null junctional zone tissues recovered from *Cited2* heterozygous (*Cited2+/-*) pregnant female rats mated with *Cited+/-* male rats and treated with either vehicle or polyI:C. CITED2 deficiency resulted in an exaggerated response of interferon-responsive transcript expression in the junctional zone (**Fig. 5E and F**).

Thus, CITED2 modulates junctional zone responses to physiological stressors, including hypoxia and a viral mimetic.

### CITED2 and human trophoblast cell development

There is supportive information connecting CITED2 to human placenta development and establishment of the EVT cell lineage. Partners in CITED2 action such as transcription factors (HIF1 and TFAP2C) and CBP/EP300 have been implicated as regulators of human trophoblast cell biology and placental disease (13, 47–49). *CITED2* expression is also downregulated in preeclamptic placental tissue (50). These relationships implicating CITED2 in human placenta pathophysiology and the above experimentation demonstrating the involvement of CITED2 in rat placentation prompted an evaluation of CITED2 in human trophoblast cell development. *CITED2* was prominently expressed in cells constituting EVT cell columns from late first trimester human placentas (12-13 weeks) except for the *CDH1 and NOTCH1* positive basal cytotrophoblast progenitor cell population (**Fig. 6A-C, SI Appendix Fig. S9**). *CITED2* transcripts co-localized with *NOTUM* (**Fig. 6D-F**), an EVT cell-enriched transcript (51), which was most prevalent in the distal region of the column (**Fig. 6E**). *CITED2* expression was also prominent in cells within a transition zone immediately proximal to the basal cytotrophoblast layer (**Fig. 6A-C**). These transition zone cells were negative for *NOTUM* (**Fig. D-F**). This places CITED2 in column locations critical for activation of the EVT cell differentiation program. The EVT cell column is a structure homologous to the junctional zone of the rat placentation site (**Fig. 1A**). EVT cell development can be effectively modeled in human TS cells (52). *CITED2* expression was significantly upregulated in EVT cells when compared to human TS cells maintained in the stem state, similar to the upregulation of *FLT4* and *NOTUM* (**Fig. 6G**). In contrast, CITED2 transcripts decline following induction of syncytiotrophoblast differentiation (***SI Appendix* Fig. S10A**). Expression of *CITED1*, a paralog of CITED2 with some similar actions (10), in human TS cells was very low and declined following EVT cell differentiation (***SI Appendix* Fig. S10B**). We next utilized shRNA mediated CITED2 silencing to investigate the potential contributions of CITED2 to the regulation of EVT cell differentiation. Differentiation involves repression of transcripts associated with the stem cell state and activation of transcripts associated with the EVT cell state. CITED2 disruption effectively interfered with the expression of CITED2 (**Fig. 6H, *SI Appendix* Fig. S11A**) and was accompanied by morphologic impairments in EVT cell-specific elongation and instead, the presence of tightly packed cell colonies resembling the TS cell stem state (**Fig. 6I**). CITED2 knockdown also inhibited the movement of trophoblast cells through Matrigel-coated transwells, an in vitro measure of cell invasion (**Fig. 6J**). RNA-seq analysis of control and CITED2 knockdown cells indicated that CITED2 possesses roles in both acquisition of the EVT state and repression of the stem state (**Fig. 6K, *SI Appendix* Fig. S11B and C**). These observations were further confirmed by RT-qPCR measurement of transcripts associated with the EVT state (*FLT4* and *NOTUM*) and stem state (*NPPB* and *PEG10*) (**Fig. 6L**). Thus, CITED2 contributes to both repression of the TS cell stem state and acquisition of the EVT cell specific developmental program.

**Figure 6.**
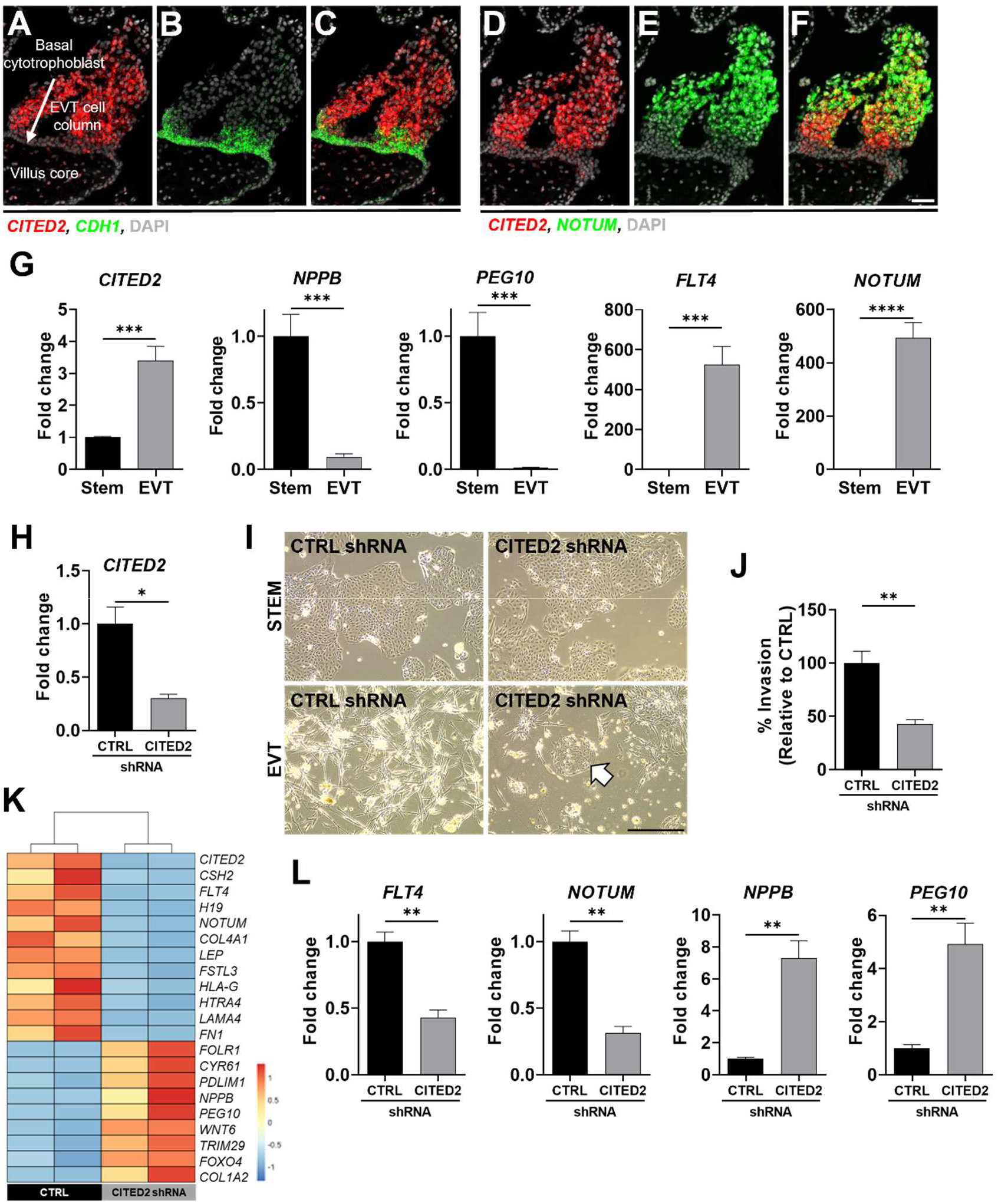
CITED2 regulates human extravillous trophoblast (EVT) cell differentiation. **A-F**. *In situ* hybridization showing *CITED2* transcript localization in first trimester (12 weeks) human placenta: C*ITED2* (**A**, red), *CDH1* (**B**, green; marker of basal cytotrophoblast), *CITED2* and *CDH1* colocalization (**C**), C*ITED2* (**D**, red), *NOTUM* (**E**, green; marker of EVT cells), CITED2 and *NOTUM* colocalization (**F**). Scale bar=50 μm. **G**. Relative expression of stem state cell signature transcripts (*NPPB, PEG10*), EVT cell signature transcripts (*FLT4* and *NOTUM*), and *CITED2* transcript in human TS cells in the stem state and following eight days of EVT cell differentiation, n=5. **H**. Relative expression of *CITED2* transcript levels in EVT cells expressing control (**CTRL**) or CITED2 shRNAs, n=3. **I**. Phase-contrast images depicting cell morphology of stem and EVT differentiated cells expressing CTRL or CITED2 shRNAs. White arrow is indicating a cluster of cells exhibiting stem-like morphology. Scale bar=500 μm. **J**. Movement of human TS cells through a Martigel coated transwell insert for cells expressing CTRL or CITED2 shRNAs. **K**. Heatmap showing select transcripts from RNA-seq analysis of human TS cells exposed to CTRL shRNA versus CITED2 shRNA during EVT cell differentiation. **L**. Relative expression of EVT cell signature transcripts (*FLT4* and *NOTUM*) and stem state cell signature transcripts (*NPPB, PEG10*) from human TS cells exposed to CTRL shRNA versus CITED2 shRNA during EVT cell differentiation, n=3. Graphs represent mean values ± SEM, unpaired t-test, *p<0.05, **p <0.01, ***p <0.001, and ****p <0.0001.

In summary, the data support the involvement of CITED2 in both rat and human deep placentation.

## Discussion

Placentation provides a means for the fetus to grow and develop in the female reproductive tract (1, 2). The structure of the mammalian placenta exhibits elements of species specificity (53, 54). However, it is also evident that there are conserved features associated with the regulation of placental development and placental function (3, 53). In this report, we provide data supporting a conserved role for CITED2 in the regulation of deep placentation in the rat and in the human.

CITED2 regulates events influencing development of the junctional zone compartment of the rat placenta and EVT cell column of the human placenta and their cell derivatives, invasive trophoblast cells and EVT cells, respectively. Most interestingly, CITED2 also contributes to regulating the plasticity of the placenta and its responses to physiological stressors.

The mouse and rat *Cited2* null phenotypes showed elements of conservation but also unique features. Prenatal lethality, adrenal gland agenesis, and exencephaly are hallmarks of the *Cited2* null mouse (15, 16, 24), but were not observed in the *Cited2* null rat. Distinct features of mouse and rat *Cited2* null phenotypes may be attributable to species differences in the role of CITED2 in embryonic development or more specifically, species differences in roles for the transcription factors that CITED2 modulates. At this juncture, evidence is not available to determine whether CITED2 biology in the mouse or rat better reflects CITED2 biology in other species, including the human. The absence of conservation between the mouse and rat should be cautionary regarding extrapolating findings with mutant rodent models to human pathophysiology, especially without additional supportive information. In contrast, the placenta represented a conserved target for CITED2 action in the mouse, rat, and human. The prominent expression of CITED2 in the junctional zone and EVT cell column directed our attention to investigating a role for CITED2 in the biology of these placental structures and their derivatives, invasive trophoblast/EVT cells. In the mouse, CITED2 also contributes to the regulation of the junctional zone and its derivatives (22). Based on widespread expression of beta-galactosidase (**lacZ**) throughout the mouse placenta in a *Cited2-lacZ* knock-in mouse model (22), CITED2 function was also investigated in the labyrinth zone (23). Cell specific trophoblast *Cited2* disruption supported a role for CITED2 in trophoblast cell-capillary patterning and transport (23). Although a growth restricted labyrinth zone was noted in the *Cited2* null rat, the absence, or potentially low levels, of CITED2 in the labyrinth zone implied that CITED2 was acting on the labyrinth zone in a non-cell autonomous role. The actions of CITED2 in the mouse labyrinth zone may represent another example of species-specificity of CITED2 action and could be responsible for the prenatal lethality observed in the *Cited2* null mouse.

Current evidence indicates that CITED2 acts as a co-regulator, modulating the recruitment of H3K27 acetyltransferases, CBP/EP300, to specific transcription factors controlling gene transcription (10). These actions can promote or inhibit CBP/EP300-transcription factor interactions and, thus, CITED2 can serve as a facilitator of gene activation or gene repression (15, 16). TFAP2C, peroxisome proliferator activating receptor gamma (**PPARG**) and HIF1 are transcription factors affected by CITED2 (15, 16, 55) with known actions on trophoblast cell development and placentation (18, 20, 21, 56). The data are consistent with CITED2 promoting placental development through stimulating transcriptional activities of transcription factors, such as TFAP2C and PPARG, and dampening placental responses to physiological stressors. CITED2 may modulate placental adaptations to hypoxia through its established role in restraining HIF-mediated transcription (15). Linkages between CITED2 and transcription factors implicated in placentation with responses to poly I:C are not evident. However, CITED2 is an established inhibitor of immune/inflammatory responses mediated by nuclear factor kappa B (45, 57) and signal transducer activator of transcription 1/interferon regulatory factor 1 (46), which could contribute to placental responses to the viral mimetic, polyI:C. CITED2 should also be viewed as an access point to other CBP/EP300-transcription factor interactions not previously known to contribute to the regulation of placentation or adaptations to physiological stressors. It is also important to appreciate that CITED2 is likely not acting as an on/off switch but instead may function as a rheostat to modulate the range of actions of specific transcription factor-CBP/EP300 interactions. CITED2 is a member of a family of transcriptional co-regulators that in mammals also includes CITED1 and CITED4 (10). CITED1 is a known regulator of placentation in the mouse (58).

Connections between CITED4 and placentation have not been described. Expression profiles of CITED1 and CITED2 in the mouse placenta overlap (22, 23, 58), as does their interaction with CBP/EP300 and transcription factors (10). Germline disruption of either *Cited1 or Cited2* in the mouse results in abnormal placentation, with altered junctional zone morphogenesis (22, 23, 58). However, their impacts on junctional zone development differ. *Cited1* null placentas showed an expansion of the junctional zone (58), whereas *Cited2* null placentas exhibited junctional zone growth restriction (22). Thus, how CITED1 and CITED2 cooperate to promote normal junctional zone morphogenesis is yet to be determined. We also presented evidence for the involvement of CITED2 in human EVT cell development but have not observed significant CITED1 expression in human TS cells or their derivatives (52). Thus, there may be a species difference in the utilization of CITED family members as regulators of placental development. Rodents require two CITED family members (CITED1 and CITED2) for normal placental morphogenesis, whereas trophoblast cell lineage development in the human may only involve CITED2.

CITED2 contributes to the regulation of the invasive trophoblast/EVT cell lineage in both the rat and human. The invasive trophoblast/EVT cell lineage is critical to the establishment of a healthy hemochorial placenta (3, 4). These cells have a key role in transforming the uterus into a milieu supportive of placental and fetal development (3, 4). Their position within the uterine parenchyma and ability to adapt to physiological stressors are fundamental to a successful pregnancy.

Disruptions in development and/or functioning of the invasive trophoblast/EVT cell lineage lead to pregnancy-related diseases, including preeclampsia, intrauterine growth restriction, and pre-term birth (59). CITED2 dysregulation has been linked to preeclampsia (50). Further interrogation of the CITED2 gene regulatory network in rat and human invasive trophoblast/EVT cells will provide insights into important developmental and pathophysiologic processes affecting hemochorial placentation and pregnancy.

## Materials and Methods

### Animals

#### Holtzman Sprague-Dawley rats were purchased from Envigo

A CITED2 deficient mouse model (15) was a gift from Dr. Yu-Chung Yang of Case Western Reserve University (Cleveland, OH). The *Cited2* mutation was moved to a CD1 mouse genetic background following >10 generations of backcrossing. Animals were maintained in a 14 h light:10 h dark cycle (lights on at 0600 h) with food and water available ad libitum. Timed pregnancies were established by cohabiting female and male rats or mice. Mating was determined by the presence of a seminal plug or sperm in the vaginal lavage for the rat and the presence of a seminal plug in the vagina for the mouse and considered gd 0.5. Pseudopregnant female rats were generated by mating with vasectomized males. Detection of seminal plugs was considered day 0.5 of pseudopregnancy.

### Hypoxia exposure

Pregnant rats were placed in a ProOX P110 gas-regulated chamber (BioSpherix) to generate a hypoxic environment [10.5% (vol/vol) oxygen] from gd 6.5 to 13.5 or gd 6.5 to 18.5 as previously described (60). Pregnant rats exposed to ambient conditions [∼21% (vol/vol) oxygen] were used as controls. Animals were euthanized at the termination of the exposures and placentation sites collected.

### Poly I:C exposure

Pregnant rats were intraperitoneally injected with poly I:C (10 mg/kg body weight, P1530-25MG, Sigma-Aldrich) on gd 13.5. Animals were euthanized 6 h following injection and placentation sites collected.

The University of Kansas Medical Center (**KUMC**) Animal Care and Use Committee approved all protocols involving the use of animals.

### Tissue collection and analysis

Rats and mice were euthanized by CO2 asphyxiation at designated days of gestation. The health and viability of placentation sites and fetuses were determined. Uterine segments containing placentation sites and fetuses were frozen in dry ice-cooled heptane and stored at -80ºC until used for histological analyses. Alternatively, placentation sites were dissected. Placentas, the adjacent uterine-placental interface tissue (also referred to as the metrial gland), and fetuses were isolated as previously described (61). Placentas were weighed and dissected into placental compartments (junctional and labyrinth zones) (61) and frozen in liquid nitrogen and stored at -80ºC until used for biochemical analyses. Fetuses were assessed for viability and morphological defects, weighed, and genotyped, and sex determined by PCR (62). Tissues from E15.5 rat fetuses and postnatal day 1 (**PND1**) newborns were dissected, fixed (E15.5 fetuses: 10% neutral buffered formalin solution; PND1 newborn tissues: 4% paraformaldehyde, **PFA**, in phosphate buffered saline, pH 7.4, **PBS**) and prepared for histological/immunohistochemical analyses. Mouse E17.5 fetal tissues were fixed in 10% neutral buffered formalin solution and tissues dissected.

Paraffin-embedded human placental tissues were obtained from the Research Centre for Women”s and Children”s Health Biobank (Mount Sinai Hospital, Toronto). Tissues were deidentified and collected following consent and approved by the University of Toronto and the KUMC human research ethics review committees.

### Generation of a *Cited2* null rat model

*CRISPR*/Cas9 genome editing was utilized to generate a CITED2 deficient rat according to procedures previously described (63). E0.5 zygotes were microinjected with guide RNAs targeting the entire coding region of the *Cited2* locus and Cas9 (***SI Appendix* Fig. S1A; Table S2**). Injected embryos were transferred to pseudopregnant rats. Offspring were screened for mutations by PCR and verified by DNA sequencing. Founder rats possessing mutations within the *Cited2* locus were backcrossed to wild type rats to confirm germline transmission.

### Genotyping and fetal sex determination

Genotyping was performed using DNA extracted from tail-tip biopsies. DNA was purified with RedExtract-N-Amp tissue PCR kit (XNAT-1000RXN, Sigma-Aldrich) using directions provided by the manufacturer. For rat *Cited2* genotyping, three primers were used to distinguish between wild type and mutant *Cited2* loci (***SI Appendix* Fig. S1B**). The sequences of these primers are provided in ***SI Appendix* Table S3**. For mouse genotyping, PCR was used to detect the neomycin resistance gene, which replaced Exon 1 and part of Exon 2 of the mouse *Cited2* gene (***SI Appendix* Fig. S1D**) (15). Primer sequences for mouse *Cited2* genotyping are provided in **Table S3**. Sex of rat fetuses was determined by PCR on genomic DNA for *Kdm5c* (X chromosome) and *Kdm5d* (Y chromosome), using primers detailed in ***SI Appendix* Table S3**, as previously described (62).

### Rat blastocyst-derived TS cell culture

Wild type and *Cited2* null rat TS cells were established, maintained, and differentiated using a previously described procedure (36). Rat TS cells were maintained in Rat TS Cell Stem State Medium (RPMI-1640 culture medium (11875093, Thermo Fisher) containing 20% fetal bovine serum (**FBS**, F2442, Sigma-Aldrich), 50 μM 2-mercaptoethanol (**2ME**, M3148, Sigma-Aldrich), 1 mM sodium pyruvate (11360070, Thermo Fisher), 100 U/ml penicillin, and 100 μg/ml streptomycin (15140122, Thermo Fisher), fibroblast growth factor 4 (**FGF4**, 25 ng/ml, 100-31, PeproTech), heparin (1 μg/ml, H3149, Sigma-Aldrich), and rat embryonic fibroblast-conditioned medium (70% of the final volume)), as previously reported (36).

Proliferation was assessed in wild type and *Cited2* null TS cells using a colorimetric assay. Wild type and *Cited2* null TS cells were plated at 1000 cells/well in 96-well cell culture treated plates. After 24, 48 and 72 h, medium was removed, cells were stained with crystal violet solution (0.4 % in methanol, C-3886, Sigma-Aldrich) for 10 min, and excess stain washed. Methanol was added to each well, incubated for 20 min and absorbance measured at 570 nm. Proliferation was expressed as fold change to 24 h values.

Differentiation was induced by the removal of FGF4, heparin, and rat embryonic fibroblast-conditioned medium, and decreasing the FBS concentration to 1%. Rat TS cells were differentiated for 12 days.

### Human TS cell culture

Human TS cells used in the experimentation have been previously described (52). The cells originated from deidentified first trimester human placental tissue obtained from healthy women with signed informed consent and approval from the Ethics Committee of Tohoku University School of Medicine. Experimentation with human TS cells was approved by the KUMC Human Research Protection Program and the KUMC Human Stem Cell Research Oversite committee. Human TS cells were maintained and differentiated into EVT cells using a previously described procedure (52). Detailed culture conditions are provided in the ***SI Appendix***.

### shRNA constructs and production of lentiviral particles

Generation of shRNA-mediated *loss-of-function* models are described in the ***SI Appendix***. shRNA sequences are included in ***SI Appendix* Table S4**.

### In vivo lentiviral transduction

Rat embryos were transduced with lentiviral particles as previously described (35). Lentiviral vector titers were determined by measurement of p24 Gag antigen by an enzyme-linked immunosorbent assay (Advanced Bioscience Laboratories). Briefly, blastocysts collected on gd

4.5 were incubated in Acid Tyrode”s Solution (EMD-Millipore) to remove zonae pellucidae and incubated with concentrated lentiviral particles (750 ng of p24/mL) for 4.5 h. Transduced blastocysts were transferred to uteri of day 3.5 pseudopregnant rats for subsequent evaluation of *Cited2* knockdown and placentation site phenotypes (**Fig. 2D**).

### In vitro lentiviral transduction

Human TS cells were plated at 50,000 cells per well in 6-well tissue culture-treated plates coated with 5 μg/mL collagen IV and incubated for 24 h. Lenti-X 293T cells were plated at 300,000 cells per well in 6-well tissue culture-treated plates coated with poly-L-lysine solution in PBS (0.001 %). Just before transduction, medium was changed, and cells were incubated with 2.5 μg/mL polybrene for 30 min at 37 °C. Immediately following polybrene incubation, cells were transduced with 500 μL of lentiviral particle containing supernatant and then incubated overnight. On the next day, medium was changed, and cells were allowed to recover for 24 h. Cells were selected with puromycin dihydrochloride (5 μg/mL, A11138-03, Thermo Fisher) for two days.

### Transient transfection

Lenti-X 293T cells were transiently transfected with a CITED2 (NM_001168388) human c-Myc and DYKDDDDK (**DDK**) tagged open reading frame clone (RC229801, Origene) using Attractene in DMEM medium supplemented with 100 U/ml penicillin, and 100 μg/ml streptomycin. On the next day, medium was replaced with fresh DMEM medium supplemented with 10% FBS, 100 U/ml penicillin, and 100 μg/ml streptomycin. Lysates were collected 48 h post-transfection.

### Matrigel invasion assay

Cell migration through extracellular matrices was assessed using Matrigel-coated transwells. A description of the invasion assay is provided in the ***SI Appendix***.

### Histological and immunohistochemical analyses

Wild type and *Cited2* null fetal, postnatal, and placental tissues were utilized for histological and immunohistochemical analyses. Protocols for assessing rat fetal development and placentation sites are presented in the ***SI Appendix***.

### *In situ* hybridization

*Cited2* transcripts in rat and human placental tissues were detected by in situ hybridization using the RNAscope® Multiplex Fluorescent Reagent Kit version 2 (Advanced Cell Diagnostics), according to the manufacturer’s instructions. Probes were prepared to detect rat *Cited2* (461431, NM_053698.2, target region: 2-1715), rat *Ceacam9* (1166771, NM_053919.2, target region: 130-847), rat *Prl7b1* (860181, NM_153738.1, target region: 28-900), human *CITED2* (454641, NM_006079.4, target region: 215-1771), human *CDH1* (311091, NM_004360.3, target region: 263-1255), human *NOTCH1* (311861, NM_017617.3, target region: 1260-2627), and human *NOTUM* (430311, NM_178493.5, target region: 259-814). Images were captured on Nikon 80i or 90i upright microscopes (Nikon) with Photometrics CoolSNAP-ES monochrome cameras (Roper).

### Western blotting

CITED2 protein in rat junctional zone tissue and DYKDDDDK (**DDK**)-tagged CITED2 protein from transfected Lenti-X 293T cells were assessed by western blotting. Information about the procedures is provided in the ***SI Appendix***.

### RT-qPCR

Total RNA was extracted by homogenizing tissues in TRIzol (15596018, Thermo Fisher), according to the manufacturer”s instructions. Purified RNA (1 μg) was used for reverse transcription using the High-Capacity cDNA Reverse Transcription kit (4368814, Applied Biosystems). Complementary DNA (**cDNA**) was diluted 1:10 and subjected to qPCR using PowerSYBR Green PCR Master Mix (4367659, Thermo Fisher), primers listed in ***SI Appendix* Table S5**, and the QuantStudio 5 Real Time PCR system (Thermo Fisher). Cycling conditions were as follows: an initial holding step (50°C for 2 min, 95°C for 10 min), followed by 40 cycles of two-step PCR (95°C for 15 s, 60°C for 1 min), and then a dissociation step (95°C for 15 s, 60°C for 1 min, and a sequential increase to 95°C for 15 s). Relative mRNA expression was calculated using the Δ Δ Ct method. Glyceraldehyde 3-phosphate dehydrogenase (*Gapdh*) was used as a reference RNA for rat samples and *POLR2A* was used for human samples.

### RNA-seq

Tissue from junctional zone compartments of gd 14.5 wild type and *Cited2* null placentas and control and *CITED2* knockdown human TS cells were collected and processed for RNA-seq analysis. RNA was extracted using TRIzol, according to the manufacturer”s instructions. cDNA libraries were prepared with Illumina TruSeq RNA sample preparation kits (RS-122-2002, Illumina). RNA integrity was assessed using an Agilent 2100 Bioanalyzer (cutoff value of RIN 8 or higher; Agilent Technologies). cDNA libraries were clustered onto a TruSeq paired-end flow cell, and sequenced (100 bp paired-end reads) using a TruSeq 200 cycle SBS kit (Illumina). Samples were run on an Illumina HiSeq2000 sequencer (tissue specimens) or Illumina NovaSeq 6000 (cells) located at the KUMC Genome Sequencing facility and sequenced in parallel with other samples to ensure the data generated for each run were accurately calibrated during data analysis. Reads from *.fastq files were mapped to the rat reference genome (*Rattus norvegicus* reference genome Rnor_6.0) or the human reference genome (*Homo sapiens* reference genome GRCh37) using CLC Genomics Workbench 20.0.4 (Qiagen). Only reads with <2 mismatches and minimum length and a similarity fraction of 0.8 were mapped to the reference genome. The mRNA abundance was expressed in reads per kilobase of exon per million reads mapped (**RPKM**). A p-value of 0.05 was used as a cutoff for significant differential expression. Functional patterns of transcript expression were further analyzed using Metascape (64).

### scRNA-seq

Uterine-placental interface tissues were dissected from gd 18.5 placentation sites (61), minced into small pieces, and enzymatically digested into a cell suspension for scRNA-seq as previously described (40, 63). Samples were then processed using Chromium Single Cell RNA-seq (10X Genomics) and libraries prepared using the Chromium Single Cell 3” kit (10x Genomics). Library preparation and DNA sequencing using a NovaSeq 6000 sequencer (Ilumina) were performed by the KUMC Genome Sequencing facility. scRNA-seq data analysis was performed as previously described (63). Briefly, the RNA sequencing data was initially processed and analyzed using the Cell Ranger pipeline. The Seurat data pipeline (version 3.1.5) was used for additional data analysis, including identification of differentially expressed genes using *FindMarkers* (65).

### Statistical analysis

Statistical analyses were performed with GraphPad Prism 9 software. Statistical comparisons were evaluated using Student”s *t* test or one-way analysis of variance with Tukey”s post hoc test as appropriate. Statistical significance was determined as p<0.05.

## Data and materials availability

All raw and processed sequencing data generated in this study have been submitted to the NCBI Gene Expression Omnibus (**GEO**) under the following accession number GSE202339. The CITED2 mutant rat model is available through the Rat Resource and Research Center (Columbia, MO).

## Acknowledgments

We thank Dr. Yu-Chung Yang of Case Western Reserve University (Cleveland, OH) for providing the *Cited2* mutant mouse model. We also thank Stacy Oxley and Brandi Miller for administrative assistance.

