## Supplementary material for "CITED2 is a Conserved Regulator of the Uterine-Placental Interface": SI Appendix

**Supporting Information for**
### CITED2 is a Conserved Regulator of the Uterine-Placental Interface

Marija Kuna,[1¶*] Pramod Dhakal,[1¶†] Khursheed Iqbal,[1] Esteban M. Dominguez,[1] Lindsey N. Kent,[1 ‡] Masanaga Muto,[1¥] Ayelen Moreno-Irusta,[1] Keisuke Kozai,[1§] Kaela M. Varberg,[1] Hiroaki Okae,[2] Takahiro Arima,[2] Henry M. Sucov,[3] and Michael J. Soares[1,4,5*]

*Corresponding authors: Marija Kuna and Michael J. Soares
 **or**

**This PDF file includes:**

Supporting text
Figures S1 to S11
Tables S1 to S5
SI References

**Other supporting materials for this manuscript include the following:**

Datasets 1 to 5 (separate excel file)

#### Materials and Methods

##### Human TS cell culture

Human TS cells were cultured in 100 mm tissue culture dishes coated with 5 µg/mL collagen IV (CB40233, Thermo Fisher). Complete Human TS Cell Medium was used to maintain cells in the stem state (DMEM/F12 (11320033, Thermo Fisher), 100 µM 2ME, 0.2% (vol/vol) FBS (16141-061, Thermo Fisher), 50 U/ml penicillin, 50 µg/mL streptomycin, 0.3% bovine serum albumin (**BSA**, BP9704100, Thermo Fisher), 1% Insulin-Transferrin-Selenium-Ethanolamine (**ITS-X**, solution (vol/vol), Thermo Fisher), 1.5 µg/mL L-ascorbic acid (A8960, Sigma-Aldrich), 50 ng/mL epidermal growth factor (**EGF**, E9644, Sigma-Aldrich), 2 µM CHIR99021 (04-0004, Reprocell), 0.5 µM A83-01 (04-0014, Reprocell), 1 µM SB431542 (04-0010, Reprocell), 0.8 mM valproic acid (P4543, Sigma-Aldrich), and 5 µM Y27632 (04-0012-02, Reprocell). To induce EVT cell differentiation, human TS cells were plated in 6-well plates pre-coated with 1 µg/mL collagen IV at a density of 60,000 cells per well. Cells were cultured in EVT Differentiation Medium (DMEM/F12 (11320033, Thermo Fisher), 100 µM 2 mercaptoethanol (**2ME**), 50 U/ml penicillin, 50 µg/mL streptomycin, 0.3% BSA, 1% ITS-X solution, 100 ng/mL of neuregulin 1 (**NRG1**, 5218SC, Cell Signaling), 7.5 µM A83-01, 2.5 µM Y27632, 4% KnockOut Serum Replacement (**KSR**, 10828028, Thermo Fisher), and 2% Matrigel® (CB-40234, Thermo Fisher)). On day 3 of EVT cell differentiation, the medium was replaced with EVT Differentiation Medium excluding NRG1 and with a reduced Matrigel® concentration of 0.5%. On culture day 6 of EVT cell differentiation, the medium was replaced with EVT Differentiation Medium excluding NRG1 and KSR, and with a Matrigel® concentration of 0.5%. Cells were analyzed on day 8 of EVT cell differentiation. To differentiate cells into syncytiotrophoblast (ST2D and ST3D), human TS cells were plated in 6-well plates pre-coated with 2.5 µg/mL collagen IV at a density of 100,000 cells per well (ST2D) or 6 cm Petri dishes (not cell culture treated) at a density of 300,000 cells per dish (ST3D). Cells were cultured in ST2D or ST3D Differentiation Medium (DMEM/F12, 100 µM 2ME, 50 U/ml penicillin, 50 µg/mL streptomycin, 0.3% BSA, 1% ITS-X solution, 2.5 µM Y27632, 4% KSR, 2 µM forskolin (F6886, Sigma-Aldrich), and with 50 ng/ml EGF only for ST3D culture). On day 3 of syncytiotrophoblast differentiation, the medium was replaced with fresh ST2D or ST3D differentiation medium. Cells were analyzed on day 6 of ST2D or ST3D differentiation.

##### Matrigel invasion assay

The in vitro invasion assay utilized Matrigel-coated transwell inserts with 8.0 µm membrane pores. Individual permeable supports (353097, Corning) were placed in a companion plate (353504, Corning) and coated with Matrigel® (CB-40234, Thermo Fisher) at concentration of 0.25 mg/ml in coating buffer (0.01 M Tris pH 8.0, 0.7 % NaCl) for 2 h at 37°C. On day 6 of human EVT cell differentiation, cells were dissociated, and 30,000 cells seeded into the upper chamber of each transwell in EVT Differentiation Medium excluding neuregulin 1 and Knockout Serum Replacement medium**.** The lower chamber was filled with the same medium containing 20% FBS. Cells were cultured at 37°C in 5% $CO_2$. After 48 h, medium was aspirated from the upper chambers, and the transwell inserts washed with PBS and then fixed with 4% PFA for 15 min at room temperature (**RT**). After fixation, transwells were washed twice with PBS, and then stained with 0.4% crystal violet solution for 15 min at RT. Transwells were then washed three times with PBS and non-invaded cells removed from the upper chambers using a cotton swab. Transwells were air dried for 24 h before the membrane was removed, placed on a slide, and imaged using Nikon 90i upright microscope with a Nikon DS-Fi1-U2 camera. Subsequently, the stained cells from four random fields were counted to calculate the percentage of invaded cells relative to control.

##### shRNA constructs and production of lentiviral particles

For rat in vivo CITED2 knockdown, *Cited2* shRNAs were designed and subcloned into pLKO.1 by using *AgeI* and *EcoRI* restriction sites as previously described (1). For human TS cell culture, two CITED2 shRNAs were designed and subcloned independently into the pLKO.1 vector at *AgeI* and *EcoRI* restriction sites. A control shRNA that does not target any known mammalian gene, pLKO.1-shSCR (plasmid 1864), was obtained from Addgene. Lentiviral packaging vectors were obtained from Addgene and included pMDLg/pRRE (plasmid 12251), pRSVRev (plasmid 12253), and pMD2.G (plasmid 12259). Lentiviral particles were produced following transient transfection of the shRNA-pLKO.1 vector and packaging plasmids into Lenti-X 293T cells (632180, Takara Bio USA). Cells were plated on 6-well tissue culture-treated plates coated with poly-L-lysine solution in PBS (0.001 %, P4707, Sigma-Aldrich) and transiently transfected using Attractene (301005, Qiagen) in Dulbecco's Modified Eagle Medium (DMEM, 11995-065, Thermo Fisher) supplemented with 100 U/ml penicillin, and 100 µg/ml streptomycin (15140122, Thermo Fisher). Cells were maintained in DMEM supplemented with 10% FBS, 100 U/ml penicillin, and 100 µg/ml streptomycin, until 24 h prior to supernatant collection, at which time the cells were cultured in Basal Human TS Cell Medium for human cells (DMEM/F12, 100 µM 2ME, 0.2% FBS, 50 U/ml penicillin, 50 µg/mL streptomycin, 0.3%, 1% ITS-X, 1.5 µg/mL L-ascorbic, 50 ng/mL EGF). Supernatants were collected for 24 h and stored frozen at −80 °C until use.

**Histological and immunohistochemical analyses**

*Rat fetal development.* E15.5 fetal tissue was fixed in 10% neutral buffered formalin, washed in PBS, and dehydrated. Tissues were then embedded in paraffin, 4 µm serial transverse sections were prepared, mounted on slides, and stained with hematoxylin and eosin (**H&E**). Lung and adrenal gland tissues were dissected from PND1 neonatal rats, fixed in 4% paraformaldehyde, washed in PBS, dehydrated, and embedded in paraffin. Transverse 6 µm sections were prepared and mounted on slides. Lung development was assessed following H&E staining. Adrenal gland development was determined by immunostaining with antibodies to cytochrome P450 family 11 subfamily A member 1, (**CYP11A1**, 1:200) (2) and tyrosine hydroxylase (**TH**, 1:200, AB152, EMD Millipore). Antigen-antibody complexes were detected using Alexa Fluor 568-conjugated goat anti-rabbit IgG (A11011, Thermo Fisher) and Alexa Fluor 488-conjugated goat anti mouse IgG (A32723, Thermo Fisher) secondary antibodies and counterstained with 4,6-diamidino-2-phenylindole (**DAPI**, 1:50,000, D1306, Invitrogen).

*Rat placentation sites.* Rat placentation sites frozen in dry-ice cooled heptane were sectioned at 10 µm, mounted on slides, fixed in 4% PFA, and treated with $H_2O_2$ in methanol. Sections were blocked with 10% goat serum (50062Z, Thermo Fisher). For immunohistochemical analysis, slides were stained using vimentin primary antibody overnight at 4°C (1:300, sc-6260, Santa Cruz Biotechnology), goat anti mouse biotin as a secondary antibody for 45 min at RT (1:150, B9904, Sigma-Aldrich), ExtrAvidin-Peroxidase for 30 min at RT (1:100, E2886, Sigma-Aldrich), and AEC substrate (SK-4200, Vector Laboratories) to stain the tissue. They were then imaged using a Nikon SMZ1500 stereoscopic zoom microscope. For immunofluorescence analysis, sections were incubated with a pan-cytokeratin antibody (1:100 or 1:150 overnight at 4°C or for 2 h at RT, F3418, Sigma-Aldrich), perforin antibodies (1:300 overnight at 4°C, TP251, Amsbio), cleaved poly [ADP-ribose] polymerase 1 (1:800 overnight at 4°C, 94885, Cell Signaling), phospho-histone 3 (1:300 overnight at 4°C, 9701, Cell Signaling), and DAPI (1:25,000 for 10 min at RT, D1306, Invitrogen). Immunostaining was visualized using fluorescence-tagged secondary antibodies, Alexa Fluor 488-conjugated goat anti mouse IgG (A11001, Thermo Fisher), and Alexa Fluor 568-conjugated goat anti-rabbit IgG (A11011, Thermo Fisher).

Slides were mounted in Fluoromount-G (0100-01, SouthernBiotech) and imaged on Nikon 80i or 90i upright microscopes with Photometrics CoolSNAP-ES monochrome cameras (Roper).

**Western blotting**

Lysates were prepared in radioimmunoprecipitation assay buffer (sc-24948A, Santa Cruz Biotechnology). Protein concentrations were determined using the DC protein assay (5000112, Bio-Rad). Protein samples were separated using sodium dodecyl sulfate-polyacrylamide electrophoresis and transferred to polyvinylidene fluoride membranes (10600023, GE Healthcare). Membranes were subsequently blocked with 5% BSA in Tris buffered saline with 0.1% Tween 20 and probed using antibodies to CITED2 (1:500, AF5005, R&D Systems), DDK (1:1,000, 14793S, Cell Signaling Technology), and/or GAPDH (1:5,000, AM4300, Thermo Fisher). Donkey anti-sheep IgG-HRP (1:5,000, sc-2473, Santa Cruz Biotechnology), horse anti-mouse IgG-HRP (1:5,000, 7076S, Cell Signaling Technology), and goat anti-rabbit IgG-HRP (1:5,000, 7074S, Cell Signaling Technology) were used as secondary antibodies. Immunoreactive proteins were visualized using chemiluminescence (Immobilon Crescendo, WBLUR0500, EMD-Millipore).

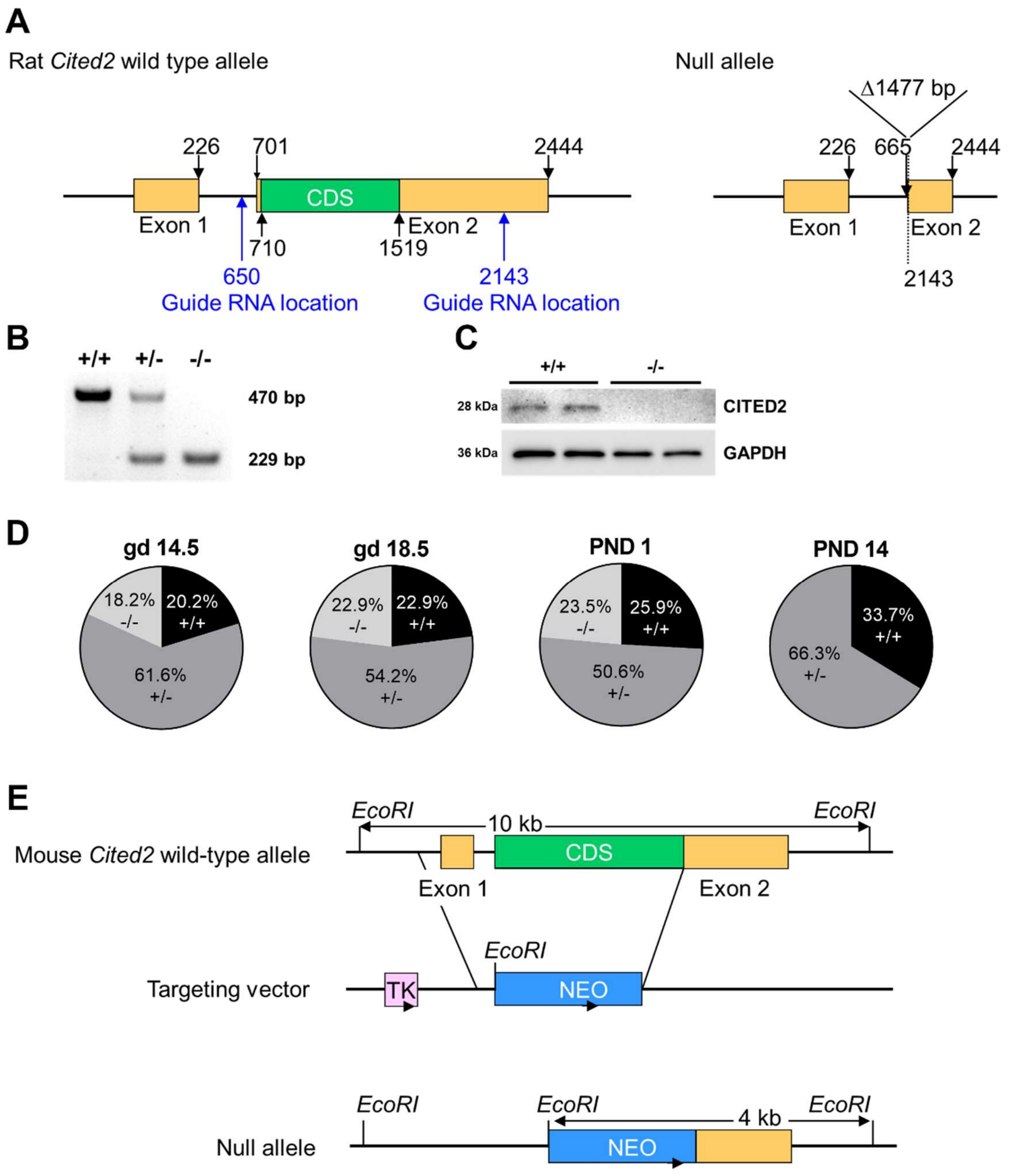


**Fig. S1. Disruption of the *Cited2* locus in the rat. A.** Schematic of the rat *Cited2* wild type and null alleles. A 1477 bp deletion was generated using CRISPR/Cas9 genome editing. The deletion included the entire coding sequence (**CDS**) of the rat *Cited*2 gene. **B**. Agarose gel electrophoresis of PCR products from fetal tail biopsies from *Cited2*$^{+/-}$ x *Cited2*$^{+/-}$ breeding showing wild type (**WT**, +/+), heterozygous (+/-) and null (-/-) genotypes. **C.** Western blot for CITED2 protein in rat junctional zone tissues from *Cited2*$^{+/-}$ x *Cited2*$^{+/-}$ breeding on gestation day (**gd**) 18.5. **D**. Pie charts depicting genotype percentages from *Cited2*$^{+/-}$ x *Cited2*$^{+/-}$ breeding on gd 14.5, gd 18.5, PND 1, and PND 14. On PND1, null (-/-) pups died within the first few h of life, n=17-61, 6-10 litters per day of gestation. **E**. Schematic representation of mouse *Cited2* wild type and mutant alleles (adapted from reference number 24). The genetic engineering resulted in deletion of the entire coding sequence (**CDS**) of the mouse *Cited2* gene.

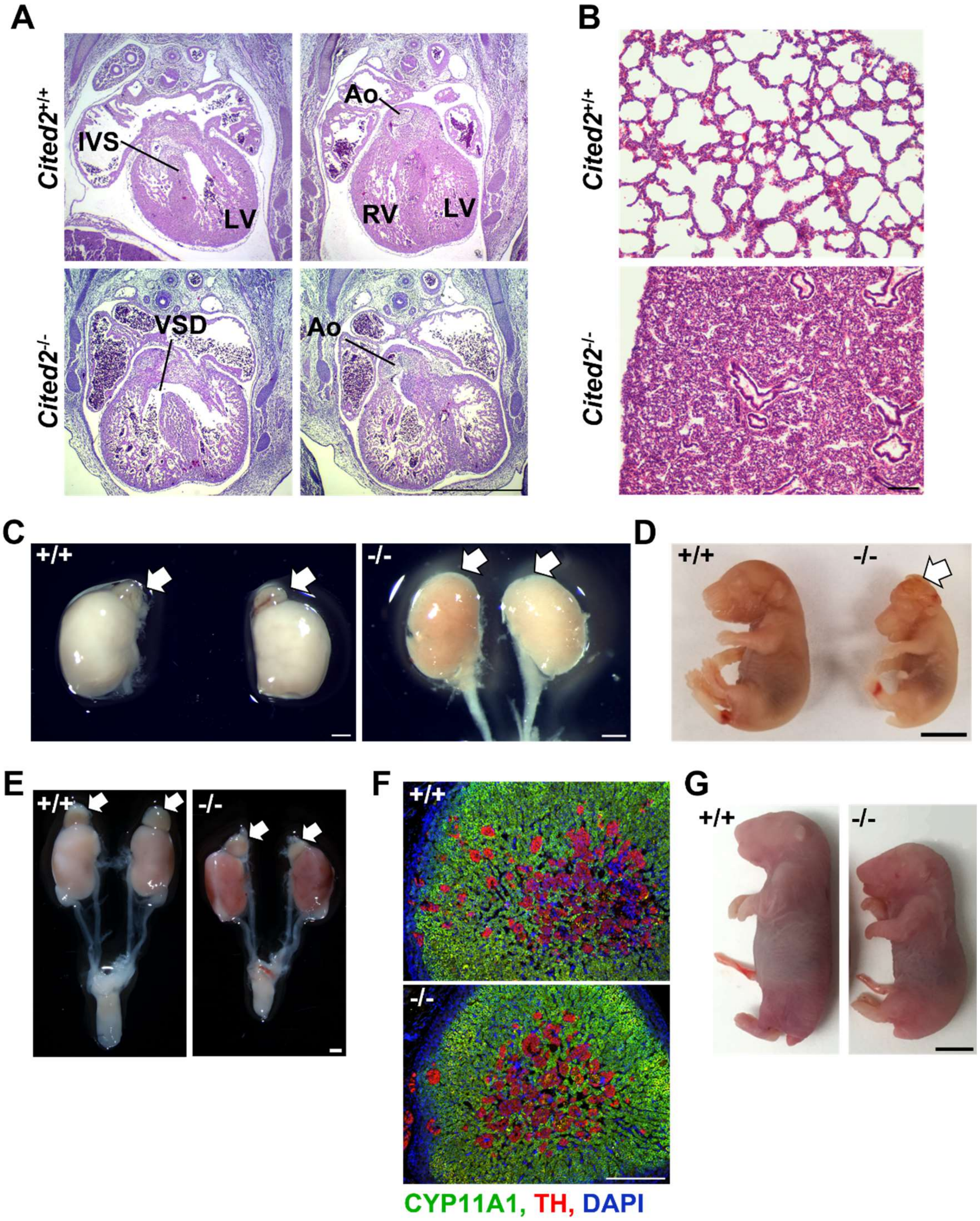


**Fig. S2. CITED2 deficiency results in death on postnatal day 1 (PND1) and heart and lung abnormalities. A.** Hematoxylin and eosin (**H&E**) stained transverse heart sections from wild type (+/+) and *Cited2* null (-/-) gestation day (**gd**) 15.5 rat embryos, scale bar=1000 µm (interventricular septum (**IVS**), left ventricle (**LV**), right ventricle (**RV**), aorta (**Ao**), ventricular septal defect (**VSD**). **B**. H&E-stained sections of lung tissue from wild type (+/+) and *Cited2* null (-/-) postnatal day 1 (**PND1**) rats, scale bar=100 µm. **C**. Kidneys with associated or missing adrenal glands (white arrows) were identified in gd 17.5 wild type (+/+) and *Cited2* null (-/-) mouse embryos, scale bar=1000 µm. **D**. Representative images for mouse gd 16.5 wild type (+/+) and

*Cited2* null (-/-) fetuses, scale bar=500 µm. The white arrow shows the exencephaly in the *Cited2* null fetal mouse. **E**. Kidneys with associated adrenal glands (white arrows) were identified in PND1 wild type (+/+) and *Cited2* null (-/-) rat pups, scale bar=1000 µm. **F**. Tissue sections from rat PND1 wild type (+/+) and *Cited2* null (-/-) adrenal glands immunostained for cytochrome P450 side chain cleavage enzyme (also called: cytochrome P450 family 11 subfamily A member 1, **CYP11A1**; green) and tyrosine hydroxylase (**TH**; red) with DAPI shown in blue, scale bar=250 µm. **G**. Representative images of wild type (+/+) and *Cited2* null (-/-) rat pups on gd 21.5, scale bar=500 µm.

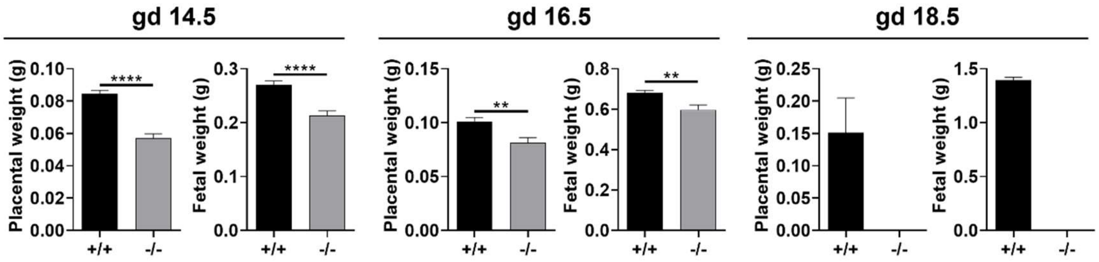


**Fig. S3. CITED2 deficiency in the mouse results in placental and fetal growth restriction.** Wild type (+/+) and *Cited2* null (-/-) placental and fetal weights from *Cited2*$^{+/-}$ x *Cited2*$^{+/-}$ breeding for the mouse on gestation day (**gd**) 14.5, 16.5, and 18.5. Graphs represent mean values ± SEM. n=18-69, 10 litters, unpaired t-test, **p<0.01, **** p<000.1.

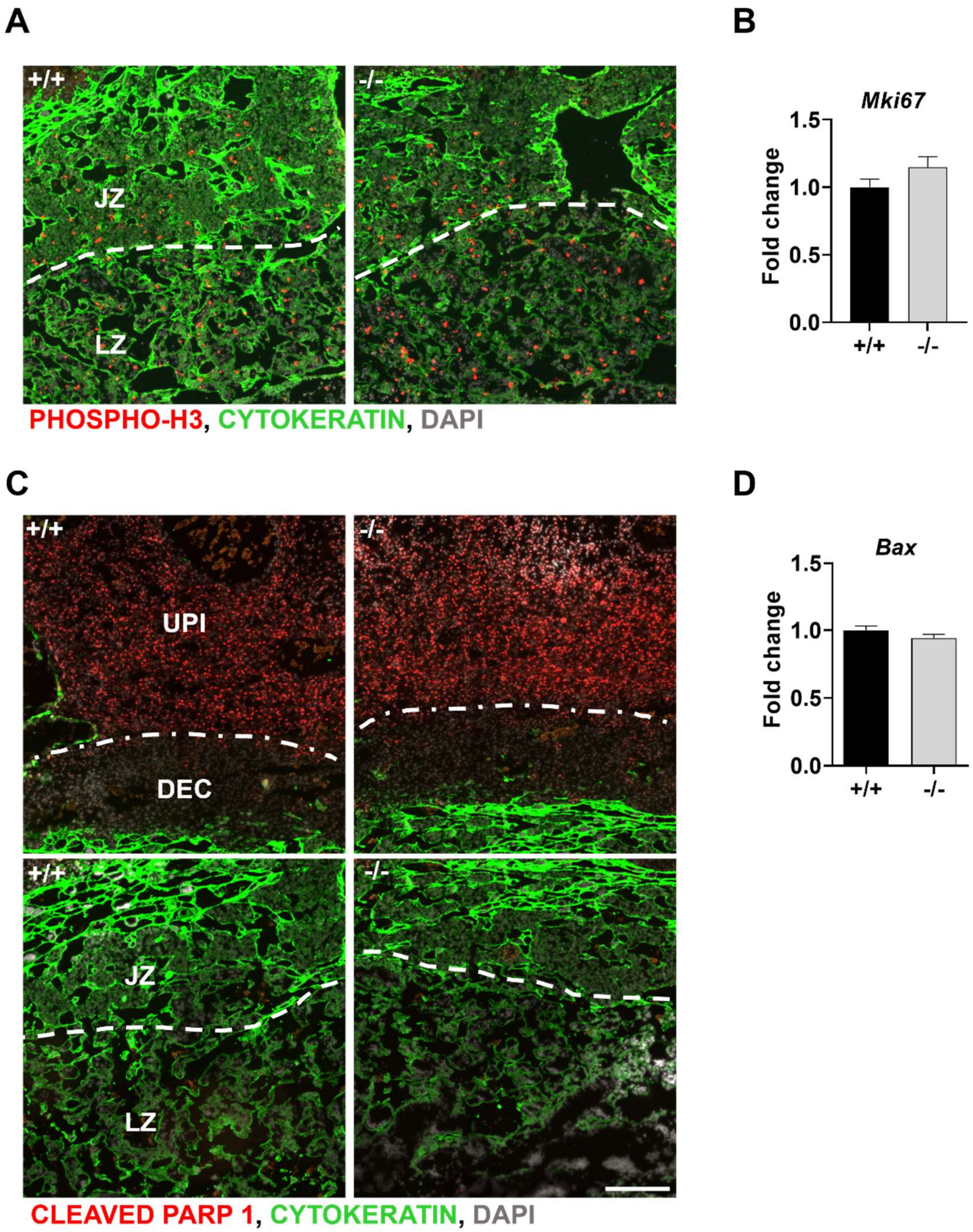


**Fig. S4. Examination of measures of proliferation and apoptosis in junctional zone tissues from wild type (+/+) and *Cited2* null (-/-) placentation sites. A.** Immunolocalization of phospho-histone 3 (**phospho-H3**) in gestation day (**gd**) 13.5 placentation sites. **B**. RT-qPCR measurements of marker of proliferation Ki-67 (***Mki67***) (proliferation associated transcript) in gd 14.5 junctional zone tissue (mean values ± SEM, n=13-21, unpaired t-test). **C**. Immunolocalization of cleaved-poly(ADP-ribose) polymerase 1 (**PARP1**) in gd 13.5 conceptus sites. Top images are from the uterine-placental interface (**UPI**), showing extensive apoptosis. Bottom images show the junctional and labyrinth zone. **D**. RT-qPCR measurement of BCL2 associated X, apoptosis regulator (***Bax)*** (apoptosis associated transcript) in gd 14.5 junctional zone tissue (mean values ± SEM, n=13-20, unpaired t-test). Cytokeratin (green) was co-localized with phospho-H3 and cleaved PARP1. DAPI (blue) staining identifies nuclei. Scale bar=250 μm.

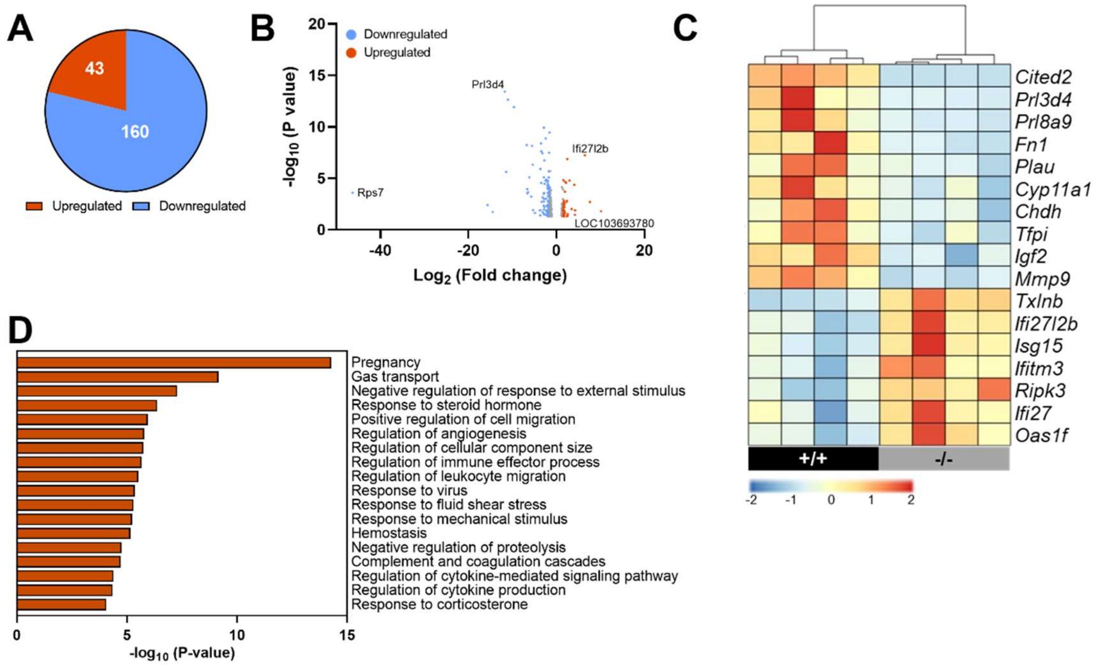


**Fig. S5. CITED2 deficiency affects the rat junctional zone transcriptome. A**. Significantly upregulated and downregulated transcripts from RNA-seq analysis of wild type (+/+) and *Cited2* null (-/-) rat gestation day (**gd**) 14.5 junctional zone tissue, n=4, $\log_2$-fold change in either direction>1.5, p<0.05. **B**. Volcano plot showing differentially expressed genes (**DEGs**) between wild type (+/+) and *Cited2* null (-/-) rat gd 14.5 junctional zone tissue samples from RNAseq analysis. *RGD1560112* and *Cited2* are outside the axis limits. **C**. Heatmap showing select transcripts from RNA-seq analysis of wild type (+/+) and *Cited2* null (-/-) rat gd 14.5 junctional zone tissue. **D**. Gene Ontology (**GO**) enriched terms for DEGs from wild type (+/+) versus *Cited2* null (-/-) rat gd 14.5 junctional zone RNA-seq analysis.

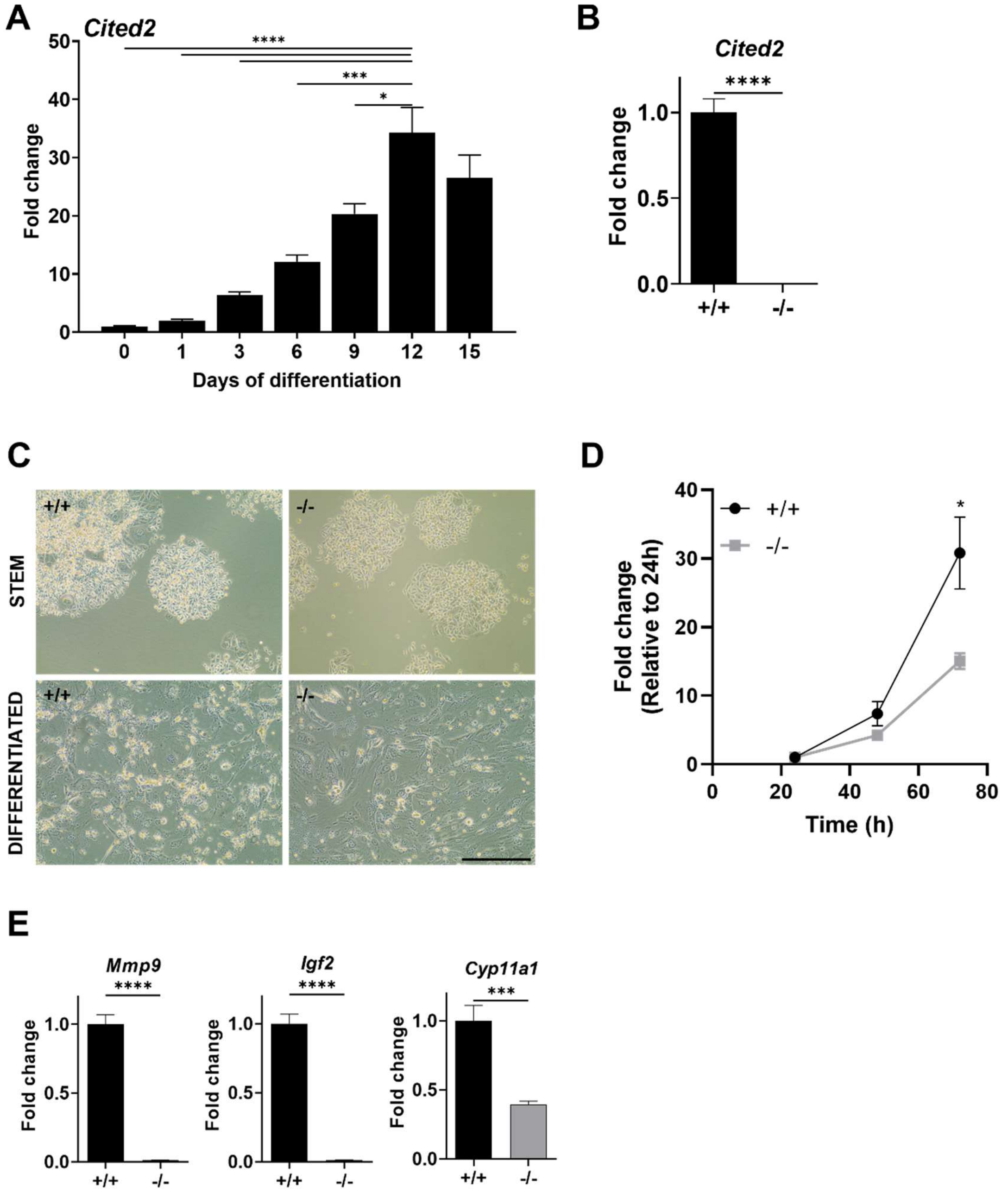


**Fig. S6. Effects of CITED2 deficiency on rat trophoblast stem (TS) cells. A**. *Cited2* transcript levels in rat TS cells maintained in the stem state and following differentiation measured by RT-qPCR (mean values ± SEM, n=4-7, one-way analysis of variance, Tukey's post-hoc test. *p <0.05, ***p <0.001, ****p <0.0001). **B**. *Cited2* transcript levels in wild type (+/+) and *Cited2* null (-/-) rat TS cells following 12 days of differentiation (mean values ± SEM, n=7-8, unpaired t-test, ****p <0.0001). **C**. Representative phase-contrast images of wild type (+/+) and *Cited2* null (-/-) rat TS cells in the stem state and following 12 days of differentiation. Scale bar=500 µm. **D**. Comparison of proliferation in wild type (+/+) and *Cited2* null (-/-) rat TS cells (mean values ± SEM, n=5-6, unpaired t-test, *p<0.05). **E**. *Mmp9, Igf2,* and *Cyp11a1* transcript levels in wild type and *Cited2* null rat TS cells following 12 days of differentiation (mean values ± SEM, n=7-8, unpaired t-test, ***p <0.001, ****p <0.0001).

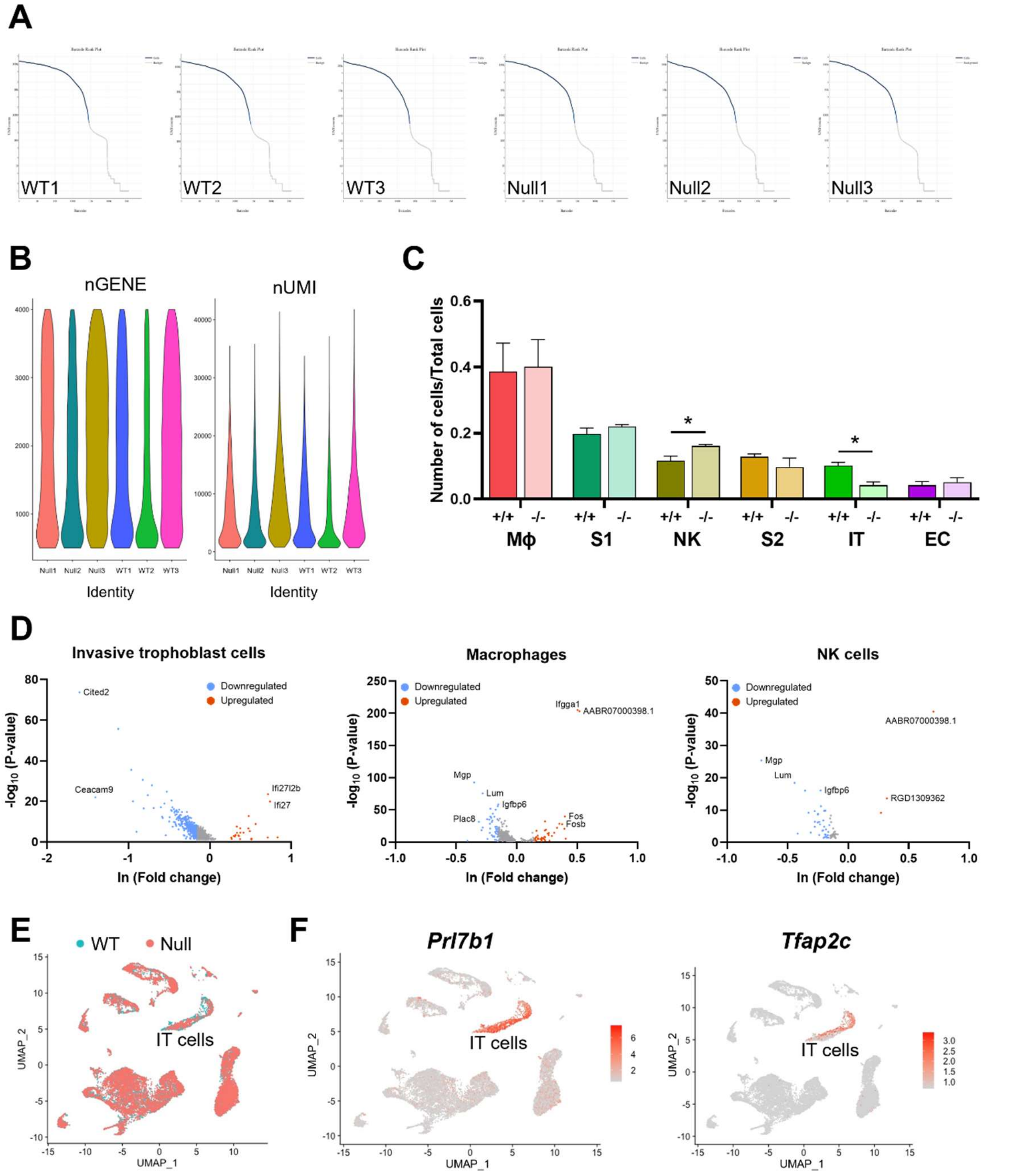


**Fig. S7. Effects of CITED2 deficiency on the uterine-placental interface.** Single cell-RNA sequencing was performed on wild type (+/+) and *Cited2* null (-/-) gestation day (**gd**) 18.5 uterine-placental interface tissue samples. **A**. Barcode rank plot generated using CellRanger software showing the distribution of barcode counts. The sharp drop denotes the UMI threshold for a barcode to be correctly categorized as a cell. **B**. Violin plot depicting the number of expressed genes (**nGene**) and number of transcripts (**nUMI**). **C**. The number of analyzed cells in respective cell clusters (mean values ± SEM, n=3, unpaired t-test, *p<0.05). Cell clusters: macrophages (**MΦ**), stromal 1 (**S1**), natural killer (**NK**) cells, stromal 2 (**S2**) cells, invasive trophoblast (**IT**) cells, and endothelial cells (**EC**). **D**. Volcano plots showing differentially expressed genes (**DEGs**) between wild type and *Cited2* null (-/-) samples for IT cell, MΦ, and NK cell clusters, p<0.05 and ln-fold change (**FC**) in either direction 0.15. **E**. UMAP plot showing cell clustering among wild type and *Cited2* null (-/-) samples. **F**. UMAP plots demonstrating *Prl7b1* and *Tfap2c* transcripts in the IT cell cluster.

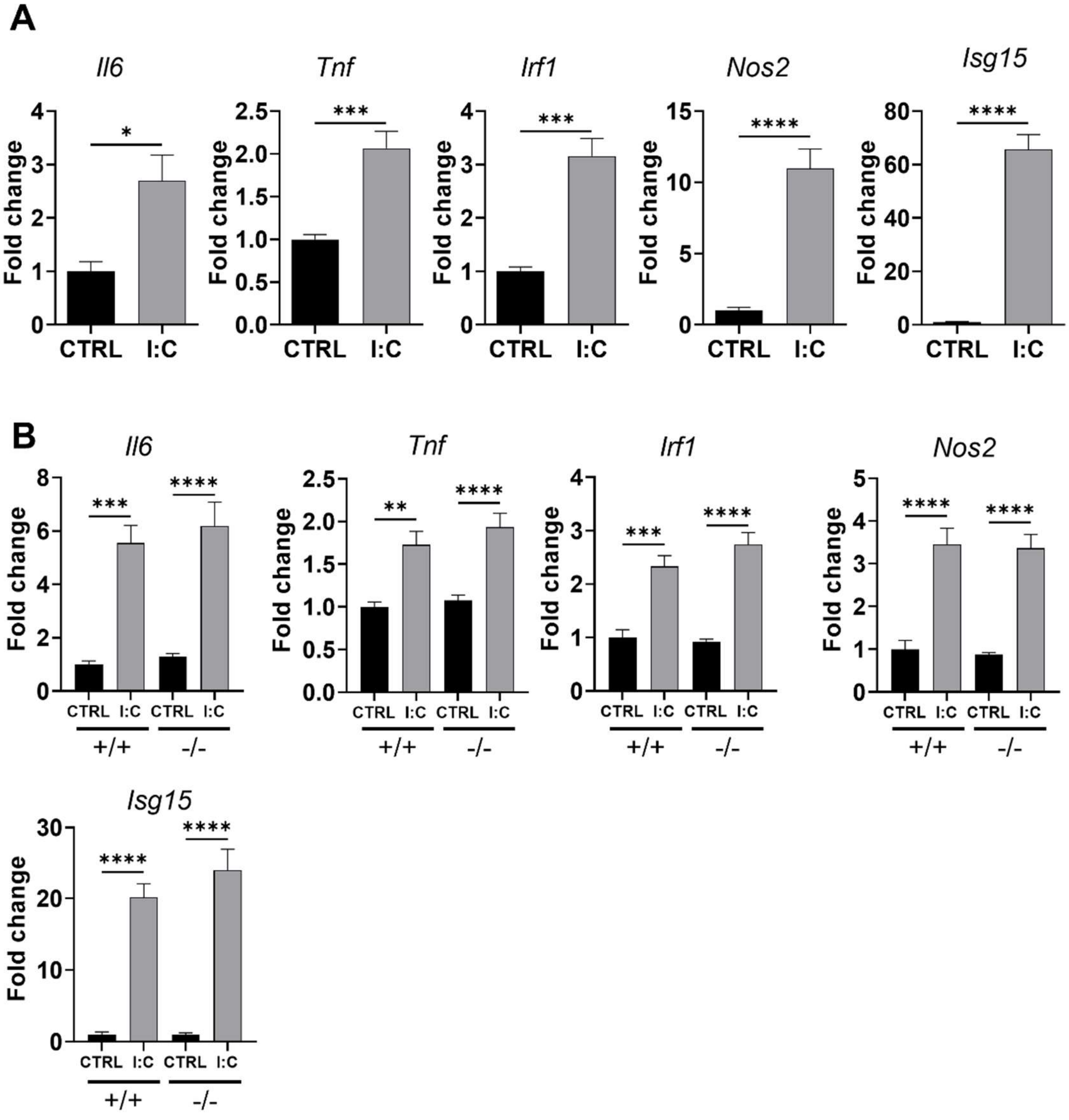


**Fig. S8. Spleen and uterine-placental interface (UPI) transcript responses to polyinosinic:polycytidylic acid (I:C) treatment.** Pregnant dams from *Cited2*$^{+/-}$ x *Cited2*$^{+/-}$ breeding on gestation day 13.5 were treated with 10 mg/kg of I:C or saline. Tissue was collected 6 h after treatment initiation. **A**. Relative expression of select transcripts in spleen tissue from saline (**CTRL**) and I:C treated animals. Graphs represent mean values ± SEM, n=5, unpaired t-test, *p<0.05, ***p <0.001, ****p <0.0001. **B**. Relative expression of select transcripts in UPI tissue from CTRL and I:C treated animals; n=8-17. Shown are mean values ± SEM, one-way analysis of variance, Tukey's post-hoc test. **p <0.01, ***p <0.001, ****p <0.0001.

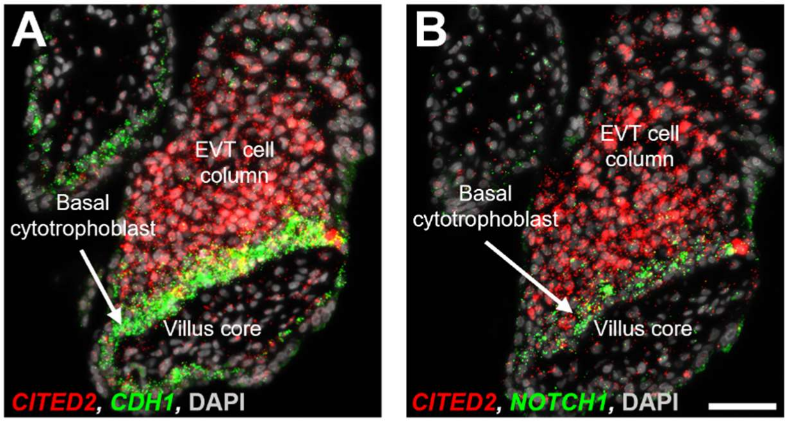


**Fig. S9. Co-localization of CITED2 expression with markers of basal cytotrophoblast of first trimester (13 weeks) EVT cell columns**. *In situ* hybridization showing co-localization of *CITED2* transcripts with *CDH1* (**A**, green; marker of basal cytotrophoblast) and *NOTCH1* (**B**, green; marker of basal cytotrophoblast). Scale bar=50 µm.

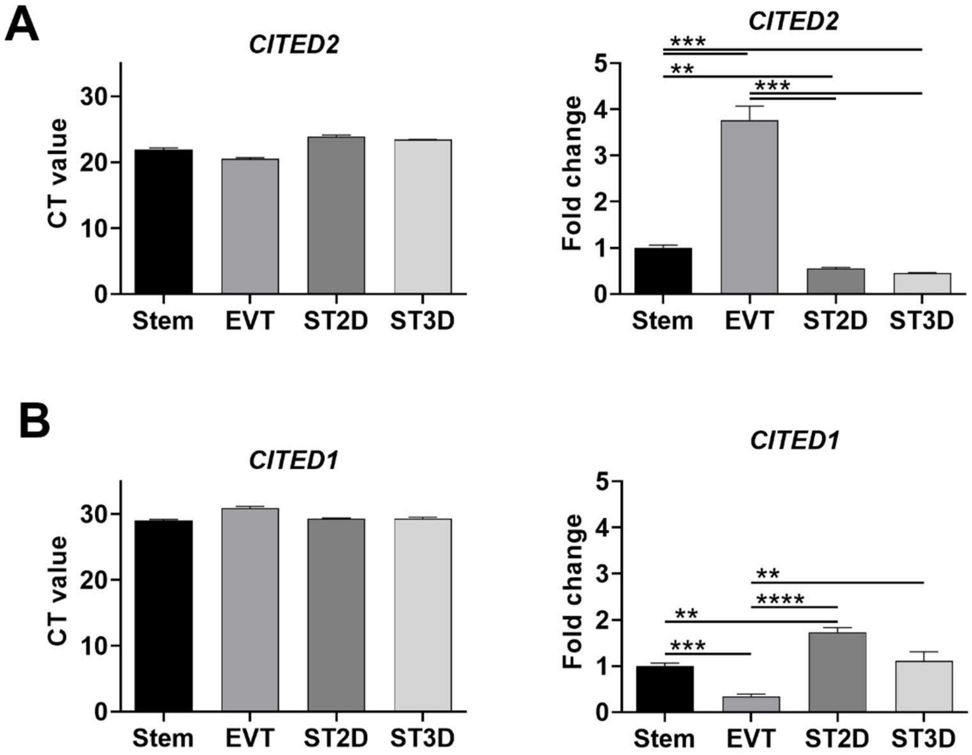


**Fig. S10. *CITED2 and CITED1* expression in human trophoblast stem (TS) cells.** RT-qPCR was performed on human TS cells in the stem state and following differentiation to extravillous trophoblast (**EVT**) cells or syncytiotrophoblast using two methods (**ST2D** and **ST3D**). **A**. Average cycle threshold (**CT**) values for *CITED2* transcripts obtained from RT-qPCR analysis (**left panel**) and relative expression of *CITED2* transcripts (**right panel**) in stem and differentiated human TS cells. **B**. Average CT values for *CITED1* transcripts obtained from RT-qPCR analysis (left panel) and relative expression of *CITED1* transcripts (right panel) in stem and differentiated human TS cells. Graphs represent mean values ± SEM, n=3-5, one-way analysis of variance and Tukey's post-hoc test, *p<0.05, **p <0.01, ****p <0.0001.

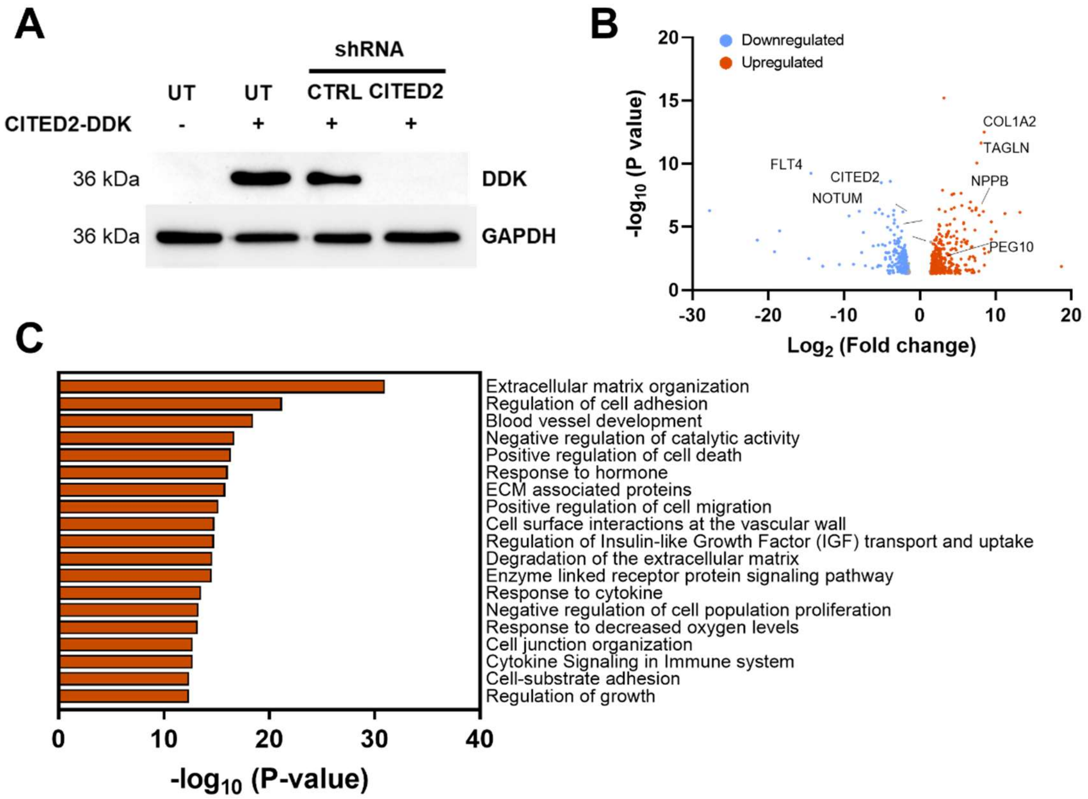


**Fig. S11. Effects of CITED2 knockdown on EVT cell differentiation. A**. Western blot for DYKDDDDK (**DDK**) in samples from Lenti-X 293T cells that were not transduced (**UT**) or transduced with control (**CTRL**) or CITED2 shRNAs and then transiently transfected with a CITED2-DDK tagged expression vector. **B**. Volcano plot showing differentially expressed genes (**DEGs**) from RNA-seq analysis of CTRL and CITED2 shRNA expressing human trophoblast stem (**TS**) cells following EVT cell differentiation. **C**. Gene Ontology enriched terms for DEGs from RNA-seq analysis of CTRL and CITED2 shRNA expressing human TS cells following EVT cell differentiation.

**Table S1.** Quality control parameters and statistics for all sample scRNA-seq libraries sequenced.

| | WT1 | WT2 | WT3 | Null1 | Null2 | Null3 |
|---|---|---|---|---|---|---|
| **Sequencing summary** | | | | | | |
| **Total # of reads** | 219,495,104 | 288,159,858 | 232,896,583 | 290,920,611 | 256,067,741 | 231,743,600 |
| **Valid Barcodes** | 98.2% | 98.2% | 97.7% | 98.3% | 97.5% | 97.6% |
| **Valid UMIs** | 100.0% | 99.9% | 99.9% | 100.0% | 99.9% | 99.9% |
| **Sequencing Saturation** | 54.8% | 63.3% | 49.5% | 51.7% | 53.1% | 50.6% |
| **Mapping details** | | | | | | |
| **Reads mapped Genome** | 95.6% | 95.0% | 93.3% | 95.6% | 94.3% | 94.4% |
| **Reads mapped confidently to Genome** | 85.2% | 84.6% | 79.5% | 84.8% | 81.4% | 80.2% |
| **Reads mapped confidently to Transcriptome** | 53.5% | 53.3% | 45.2% | 52.1% | 47.4% | 46.6% |
| **Cell Summary** | | | | | | |
| **Estimated # of cells** | 7,518 | 6,856 | 4,749 | 6,777 | 6,339 | 5,553 |
| **Fraction Reads in Cells** | 87.4% | 89.2% | 89.6% | 92.4% | 90.6% | 89.6% |
| **Total Genes Detected** | 55,799 | 17,597 | 17,029 | 17,562 | 17,105 | 17,045 |
| **Medial UMI counts/ Cells** | 6,358 | 5,974 | 17,029 | 5,543 | 3,935 | 4,745 |

**Table S2.** Guide RNA sequences.

| Sequence | Location |
|---|---|
| **GGTGGGCGACACGTGTGGCGNGG** | 650-672 |
| **GGGCCAAACCGTTCTGGATCNGG** | 2124-2143 |

**Table S3.** Genotyping primers.

| Primer | FW Sequence | RV Sequence |
|---|---|---|
| **Rat Cited2** | TGCTGGTGGGTTCTCTCTTG | CTGTGAAATGTTTGCCACTGAC |
| | CTTGACATCCACCTCCCTTATG | |
| **Rat Kdm5c** | TTTGTACGACTAGGCCCCAC | CCGCTGCCAAATTCTTTGG |
| **Rat Kdm5d** | TTGGTGAGATGGCTGATTCC | |
| **Mouse Neo** | TGCTCCTGCCGAGAAAGTATCC ATCATGGC | CGCCAAGCTCTTCAGCAATATC ACGGGTAG |
| **Mouse Cited2** | TGTTCCGAGCAGAAATCGCA | GGGGTTGCAATCTCGGAAGT |

**Table S4.** shRNA sequences for in vivo and in vitro knockdown experiments.

| Name | Target Sequence | Species |
|---|---|---|
| **shCited2-2** | CATCGACGAGGAAGTGCTTATCTC | Rat |
| **shControl** | CCTAAGGTTAAGTCGCCCTCG | Human/Rat |
| **shCITED2-7** | CATCGACGAGGAAGTTCTTAT | Human |
| **shCITED2-9** | AGCTGTTGACTCGATCGAAAC | Human |

**Table S5** Primers used for RT-qPCR.

| Primer | FW Sequence | RV Sequence |
|---|---|---|
| **Human *CITED1*** | AGGATGCCAACCAAGAGATG | TGGTTCCATTTGAGGCTACC |
| **Human *CITED2*** | GGTTTGGACCGCATCAAG | GATCGAGTCAACAGCTCACTCT |
| **Human *FLT4*** | GTGACGTGTGGTCCTTTGG | CTGGCAGAACTCCTCATTGAT |
| **Human *NOTUM*** | ACAGGGATCCTGTCCTCACA | CTCCAAACATCACTGGAGCA |
| **Human *NPPB*** | CTTTCCTGGGAGGTCGTTCC | GTTGCGCTGCTCCTGTAAC |
| **Human *PEG10*** | GGAGAACAGCGGAGAAGGTC | CAAAACCCGCTTATTTCACGC |
| **Human *POLR2A*** | TCCGTATTCGCATCATGAAC | TCATCCATCTTGTCCACCAC |
| **Rat *Bax*** | CGGCGAATTGGAGATGAACTGG | CTAGCAAAGTAGAAGAGGGCAACC |
| **Rat *Ceacam9*** | GGCGGAAACTCCGTTTTGTT | TGGTGGGTGTTAAGTACCGC |
| **Rat *Cyp11a1*** | CAGCTGCCTGGGATGTGATT | AGGACACCAGGGTACTTGCT |
| **Rat *Cited2*** | TCTTGGCTGCATGAACTTTG | GGGAGACAGCCAACTTGAAA |
| **Rat *Gapdh*** | GACATGCCGCCTGGAGAAAC | AGCCCAGGATGCCCTTTAGT |
| **Rat *Ifi27l2b*** | CAGCTCATCATGTGGGGAGAAA | GCTGCAGCAGACATCATCTTG |
| **Rat *Igf2*** | GGAGGGGAGCTTGTTGACAC | AGCAGCACTCTTCCACGATG |
| **Rat *Il6*** | CACAAGTCCGGAGAGGAGAC | CAGAATTGCCATTGCACAAC |
| **Rat *Irf1*** | TCATGCCAGACAGCACCAC | GGCTGCCACTCAGACTGTTC |
| **Rat *Isg15*** | ACAGCCATGACCTGGAACCTAAA | CTGGCACACCACTCTTCTGA |
| **Rat *Mki67*** | AGTGGCCAAACAGACTTGCT | AGGCACTCCCTCACTCTTGT |
| **Rat *Mmp9*** | TCCAGTAGACAATCCTTGCAATGTG | CTCCGTGATTCGAGAACTTCCAATA |
| **Rat *Mx2*** | GCTCATCTCACACATCTGTAAATCT | GAGCTTGGTGAATAGGCGACA |
| **Rat *Nos2*** | AGGCCACCTCGGATATCTCT | GCTTGTCTCTGGGTCCTCTG |
| **Rat *Oasl2*** | TAAGGGCGACCGGCCTATCA | GAGGTAGTGCGCGATAGATACC |
| **Rat *Prl7b1*** | CCGTCATACTGTCTCAGCACATC | AGCTGTTGAGACCATTGACAACAAA |
| **Rat *Tnf*** | GATCGGTCCCAACAAGGAG | TCCTCCGCTTGGTGGTT |

**Dataset S1 (separate file).** Differentially regulated transcripts identified in the RNA-seq analysis of rat gestation day 14.5 junctional zone tissue from wild type and *Cited2* null rats.

**Dataset S2 (separate file).** Differentially regulated transcripts identified in the invasive trophoblast cells from scRNA-seq analysis of wild type and *Cited2* null rat gestation day 18.5 uterine-placental interface tissue.

**Dataset S3 (separate file).** Differentially regulated transcripts identified in macrophages from scRNA-seq analysis of wild type and *Cited2* null rat gestation day 18.5 uterine-placental interface tissue.

**Dataset S4 (separate file).** Differentially regulated transcripts identified in NK cells from scRNA-seq analysis of wild type and *Cited2* null rat gestation 18.5 uterine-placental interface tissue.

**Dataset S5 (separate file).** Differentially regulated transcripts identified by RNA-seq analysis in stem and EVT differentiated human TS cells following *CITED2* knockdown.
